# Nanodomain-mediated lateral sorting drives polarization of the small GTPase ROP2 in the plasma membrane of root hair cells

**DOI:** 10.1101/2021.09.10.459822

**Authors:** Vanessa Aphaia Fiona Fuchs, Philipp Denninger, Milan Župunski, Yvon Jaillais, Ulrike Engel, Guido Grossmann

## Abstract

Formation of root hairs involves the targeted recruitment of the cellular growth machinery to the root hair initiation domain (RHID), a specialized site at the plasma membrane (PM) of trichoblast cells. Early determinants in RHID establishment are small GTPases of the Rho-of-plants (ROP) protein family, which are required for polarization of downstream effectors, membrane modification and targeted secretion during tip growth. It remains, however, not fully understood how ROP GTPases themselves are polarized. To investigate the mechanism underlying ROP2 recruitment, we employed Variable Angle Epifluorescence Microscopy (VAEM) and exploited mCitrine fluorophore blinking for single molecule localization, particle tracking and super-resolved imaging of the trichoblast plasma membrane. We observed the association of mCit-ROP2 within distinct membrane nanodomains, whose polar occurrence at the RHID was dependent on the presence of the RopGEF GEF3, and found a gradual, localized decrease of mCit-ROP2 protein mobility that preceded polarization. We provide evidence for a step-wise model of ROP2 polarization that involves (i) an initial non-polar recruitment to the plasma membrane via interactions with anionic phospholipids, (ii) ROP2 assembly into membrane nanodomains independent of nucleotide-binding state and, subsequently, (iii) lateral sorting into the RHID, driven by GEF3-mediated localized reduction of ROP2 mobility.

## Introduction

Formation of functional cell shape involves the establishment of one or multiple polarity axes. To this end, the underlying growth machinery is recruited to a specific site within the plasma membrane, resulting in locally restricted activation and regulation of growth. The central questions regarding the establishment of a polar domain are: How is the site for polar growth determined, how are proteins of the growth machinery specifically recruited to this site and how is such a polar domain maintained?

The formation of root hairs in Arabidopsis thaliana is well suited to address these questions and to investigate the processes underlying polar domain establishment and targeted recruitment of the growth machinery. Root hairs are tip-growing, tubular protrusions with a diameter of approximately 10 µm that emerge from specialized epidermal root cells called trichoblasts, typically 10 µm away from the basal (root meristem oriented) end of the cell (Grierson et al., 2014). In Arabidopsis, trichoblasts are organized in cell files in which development of individual root hair cells progresses with increasing distance from the root meristem, thus depicting a developmental time line. As a result, the future site of hair outgrowth is predictable and can be investigated even before visible morphological changes occur (Denninger et al., 2019).

Small Rho-type GTPases of the Rho of Plants (ROP) family are key players in root hair growth, known to regulate cytoskeletal dynamics and vesicle trafficking (Feiguelman et al., 2018). Like all small GTPases, ROPs act as molecular switches, that can be present in an active (GTP-bound) and an inactive (GDP-bound) state (Zheng and Yang, 2000). Activation and inactivation of ROPs is facilitated by proteins of the guanine nucleotide exchange factor (RopGEF) and GTPase activating protein (RopGAP) families, respectively (Feiguelman et al., 2018), as well as the only known Arabidopsis Rho GTPase GDP dissociation inhibitor (RopGDI) SUPER-CENTIPEDE 1 (SCN1) (Carol et al., 2005).

The ROP family member ROP2 is a very early determinant during Arabidopsis root hair development, polarly localizing to the root hair initiation domain (RHID) within the plasma membrane (PM) before root hair emergence is morphologically visible (Jones et al., 2002). ROP2 polarization at the RHID can be detected as early as 4 cells before root hair bulging (Fig.1) (Denninger et al., 2019). Even though the role of ROP2 during root hair growth is well studied, the mechanism leading to the accumulation of a sufficient number of ROP2 molecules, resulting in hair outgrowth, remains not fully understood. Two mechanisms seem possible: RHID-directed, targeted secretion or untargeted attachment of ROP2 to the PM and subsequent, lateral sorting into the RHID.

**Fig. 1.**
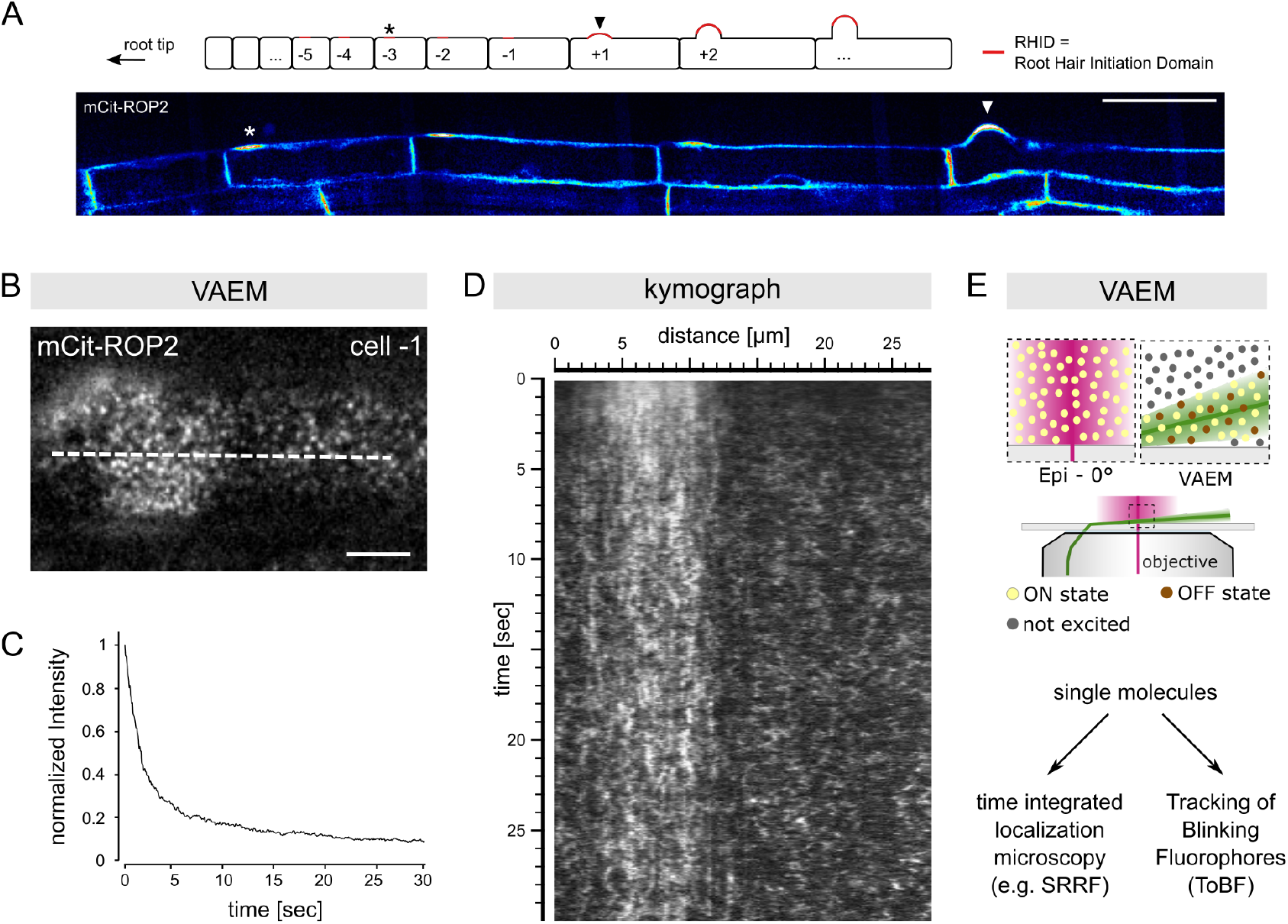
Fluorophore blinking allows to improve the spatio-temporal resolution at the plant plasma membrane. (A) schematic representation of a root hair cell file, which from root tip to shoot depicts a developmental timeline. Cell stage determination is performed by counting from the first visible bulge, which is named +1. Younger cells are assigned to negative, older cells to positive numbers. Below, a trichoblast cell file of a mCitrine (mCit)-ROP2 expressing plant line is shown. Asterisk marks the first polar accumulation of the fusion protein to the root hair initiation domain (RHID) at cell stage -3; the arrow indicates cell stage +1, where the first bulging root hair is visible. Scale bar represents 20 µm. (B) VAEM micrograph of a -1 trichoblast expressing mCit-ROP2; dashed line indicates the line along which the kymograph in (D) is drawn; scale bar represents 5 µm. (C) normalized fluorescence intensity at the RHID over the course of the time-lapse stack represented in (B). (D) kymograph of the VAEM time-lapse stack represented in (B); for better visualization of the mCit-ROP2 protein, the contrast was maximized in each time frame. (E) graphical representation of epifluorescence microscopy and VAEM; high laser power together with a highly inclined incident laser beam causes fluorophore blinking (represented by brown and yellow dots) leading to single molecule detection, which can be used for time-integrated localization microscopy (left arrow) or tracking of blinking fluorophores (ToBF; right arrow).

Recently, we could show that the RopGEF GEF3 serves as a landmark protein for the RHID and is necessary and sufficient to polarize ROP2 (Denninger et al., 2019). In addition, we have provided evidence for protein-protein interactions between the N-terminus of ROP2 and GEF3. However, whether GEF3-dependent polarization of ROP2 to the RHID involves targeted secretion or lateral sorting into the RHID, remains to be tested.

To investigate the mechanism underlying ROP2 polarization at the RHID, we employed Variable Angle Epifluorescence Microscopy (VAEM) Konopka and Bednarek (2008). Compared to conventional confocal laser scanning microscopy, VAEM increases the signal to background ratio by only illuminating fluorophores in a small sheet at the surface of the specimen with a highly inclined incident laser beam (Fig.1E). VAEM imaging can be performed in nearly real time, making it a suitable method to study the dynamic behavior of proteins in the PM of trichoblasts. In addition, we take advantage of the intrinsic blinking behavior of mCitrine, to perform single molecule localization microscopy. Time-integration of localization events of single mCit-ROP2 molecules revealed a sub-compartmentation of the RHID into GEF3-dependent ROP2 nanodomains. Additionally, we performed particle tracking of mCit-ROP2 molecules over short distances and analyzed overall ROP2 protein mobility during RHID development. We found that ROP2 mobility was reduced prior to its polarization at the RHID in a GEF3-dependent manner. We furthermore show, that differential activation of ROP2 is necessary for its polarization at the RHID. In addition, we show that electrostatic interactions between ROP2 and anionic lipids in the PM are a prerequisite for targeting ROP2 to the PM, but are not sufficient for its polarization at the RHID. Overall, we present evidence for a step-wise polarization mechanism that involves lateral sorting of ROP2 in the PM to facilitate RHID set-up.

## Results

### Fluorophore blinking allows to improve the spatio-temporal resolution at the plant plasma membrane

Citrine and related fluorescent proteins (FPs) have been reported to exhibit photochromic behavior, manifesting in fluorescence intermittency, also called fluorophore blinking (Dickson et al., 1997; Fölling et al., 2008). In this study we exploited the intrinsic feature of mCitrine to undergo fluorophore blinking, in order to characterize the dynamic processes underlying ROP2 polarization at the RHID with a high spatio-temporal resolution.

Fluorophore blinking is based on the stochastic process in which electrons of a fluorophore transition into a transient, non-emitting and non-excitable meta-state, also called the dark state. The more energy is transferred to the fluorophore, the higher the likelihood that its electrons undergo this transition (Vogelsang et al., 2010). With an increasing number of FPs entering the dark state, the density of excitable FPs decreases and consequently the likelihood to detect single FPs increases. To test for the ability of mCitrine to undergo fluorophore blinking in plant cells, we investigated the actin probe LifeAct (Riedl et al., 2008) fused to mCitrine in comparison to a LifeAct-mNeonGreen fusion, both transiently expressed in *Nicotiana benthamiana* leaves, by applying high excitation intensity. To minimize phototoxic effects and increase contrast, we employed Variable Angle Epifluorescence Microscopy (VAEM) (Konopka and Bednarek, 2008), a technique involving highly inclined illumination of the specimen. VAEM is typically performed on Total Internal Reflection Fluorescence (TIRF) microscopes, taking advantage of light refraction at the interface of two media with different refractive indices (glass – water) but uses a subcritical illumination angle. In contrast to TIRF microscopy (TIRFM), fluorophore excitation in VAEM is, therefore, not based on an evanescence wave protruding up to 100 nm from the interface into the specimen, but on direct excitation with a variable shallow angle (Konopka and Bednarek, 2008). Thus, VAEM is a suitable alternative to TIRFM for high contrast imaging in plant cells, which are encapsulated with cell walls and, at the same time, reaches the intensities required for dark state transitions of fluorophores.

VAEM time-lapse stacks revealed, that with increasing laser power, mCitrine-tagged LifeAct exhibited a punctate, “pearls on a string”-like appearance (Fig.S1A-C, supplemental movie 1). This was not the case for LifeAct tagged with mNeonGreen (Fig.S1D-F, supplemental movie 1), demonstrating that, under our imaging conditions, mCitrine was able to exhibit fluorophore blinking in plant cells and that this phenomenon was fluorophore-specific.

Next, we used VAEM on trichoblasts expressing ROP2 tagged with mCitrine at its N-terminus (mCit-ROP2, Fig.1A). mCitrine molecules transitioning into the dark state manifested as fast decrease of fluorescence intensity within the first few seconds of recording (Fig.1C). In VAEM time-lapse stacks of trichoblasts expressing mCit-ROP2, single puncta became visible that showed appearance and disappearance and little lateral movement (Fig.1B, supplemental movie 2). A kymograph, drawn along the trichoblast surface, showed the polarization of ROP2 molecules within the RHID, as indicated by differential fluorescent intensity in- and outside the RHID. In addition, mCit-ROP2 appeared as discontinuous stripes, indicating a relatively high positional stability at the PM. Outside the RHID, mCit-ROP2 appeared in a dot-like manner in the kymograph, indicating a lower protein amount and, thus, less frequent detection of individual, blinking mCit-ROP2 molecules (Fig.1D).

Together, the ability to detect single molecules enables to follow two distinct approaches (Fig.1E): On the one hand, the detection of single blinking FPs allows to integrate their position over time, allowing for the computation of a super-resolved image. On the other hand, it is possible to perform particle tracking of single blinking molecules, which, in comparison to methods analyzing a small number of single particles (e.g. single particle tracking) or bulk protein behavior (e.g. Fluorescence Recovery After Photobleaching - FRAP), provides a detailed picture of the diverse range of protein mobilities at the plasma membrane. The resulting large numbers that are obtained per cell and experiment offer the prospect of revealing even subtle differences in lateral mobility in protein subpopulations that may be sufficient to break symmetry and mark the beginning of cell polarization.

### ROP2 partitions in nanodomains within the RHID

To investigate the process of ROP2 polarization during RHID development, we next aimed to perform single molecule localization (SML) in mCit-ROP2 expressing plants. For the time-integrated localization of individual mCit-ROP2 molecules from VAEM data, we applied the Super Resolved Radial Fluctuation (SRRF) algorithm (Gustafsson et al., 2016). SRRF does not only allow to increase spatial resolution by SML, but also emphasizes objects with high positional stability, which in our case reflects the re-occurrence of blinking events at a specific location (the brighter the pixel appears, the more often a blink event has been localized here over time). Additionally, the SRRF algorithm performs sub-pixel localization prior to time integration, decreasing the impact of movement within the sample (Gustafsson et al., 2016).

SRRF reconstructions of mCit-ROP2 showed, that the fusion protein accumulated in dots with high positional stability (representing a reduction in lateral mobility) evenly distributed across the cell surface in cell stage -4 (Fig.2A). We were able to confirm similar sub-compartmentation of the RHID into mCit-ROP2 nanodomains, using high-resolution confocal laser scanning microscopy (CLSM) (Fig.S2A), however, lateral resolution was low due to motion blur and additionally, fast bleaching of the sample only allowed the acquisition of a single time-point. Therefore, detailed analysis of mCit-ROP2 nanodomains localization and protein mobility, was performed using VAEM and SRRF.

**Fig. 2.**
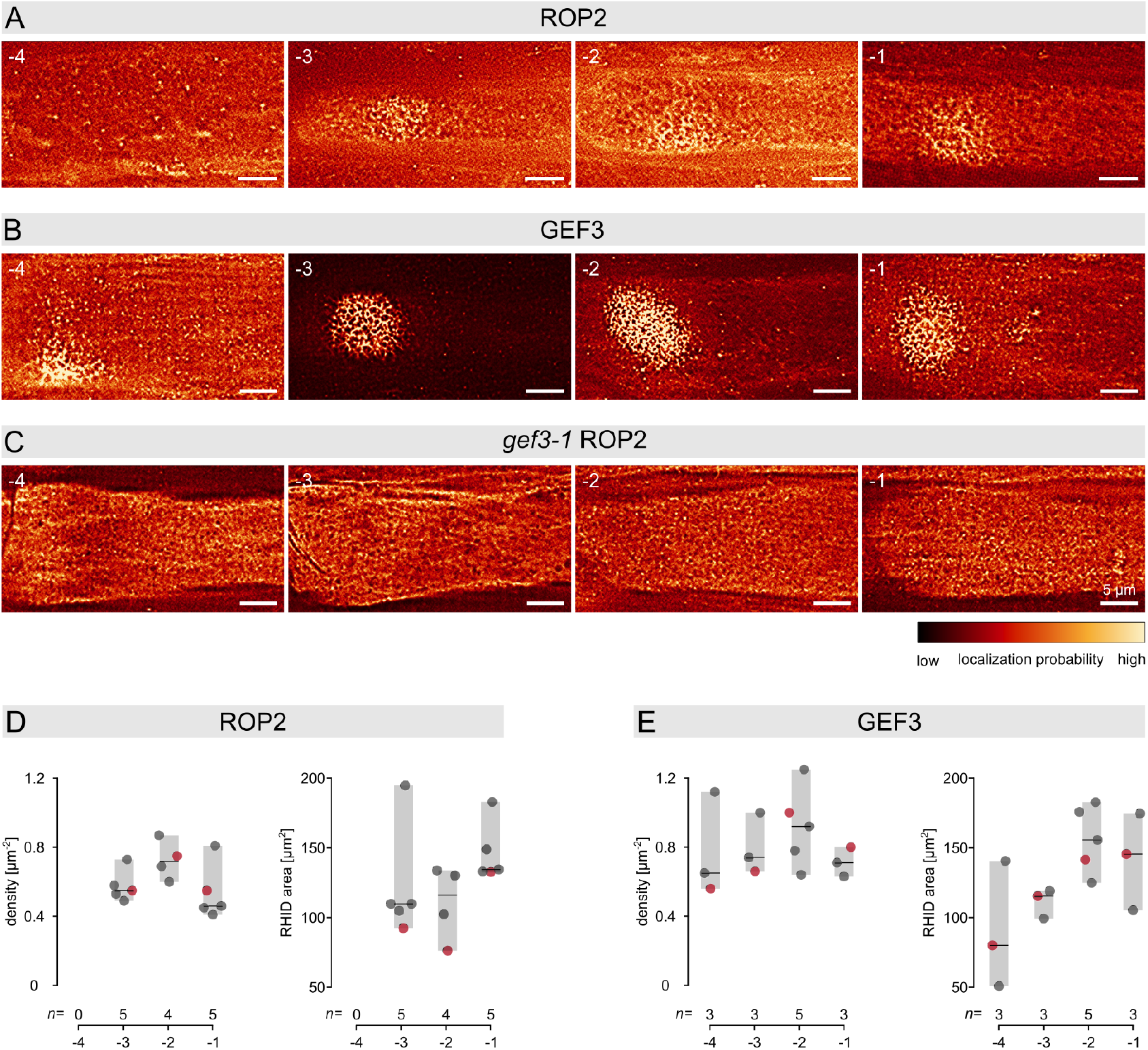
ROP2 is recruited into stable nanodomains at the RHID in a GEF3-dependent manner. (A-C) Super Resolution Radial Fluctuation (SRRF) reconstructions of VAEM micrographs of ROP2 (A), GEF3 (B) and ROP2 in the *gef3-1* mutant background (C) in different developmental cell stages (−4 to -1). The root tip is always located to the left side of the cells. Stability of the structures is false-colored as indicated; scale bars represent 5 µm. (D, E) Quantification of the density of nanodomains and the area of the RHID for ROP2 (D) and GEF3 (E). Centre lines represent median values; gray boxes indicate the data range; n represents the number of cells measured. Red data points represent the analysis of the above shown images.

Over the course of cell differentiation, ROP2 nanodomains were observed predominantly at the RHID. Outside of the RHID, the overall appearance of mCit-ROP2 was less positionally stable, indicated by comparably low pixel values depicted in the representation obtained by the SRRF analysis. Overall, SRRF reconstructions of mCit-ROP2 revealed that its distribution at the PM of the RHID was not homogenous but compartmentalized into nanodomains. The density of immobile nanodomains at the RHID remained constant within the same developmental time frame, while the area of the RHID minimally increased over time (Fig.2D).

### The presence of ROP2 nanodomains at the RHID is dependent on GEF3

To investigate whether the occurrence of ROP2 in nanodomains at the RHID might be dependent on GEF3, which we have shown earlier to be necessary and sufficient to polarize ROP2 at the RHID (Denninger et al., 2019), we next used VAEM on plant lines expressing mCit-GEF3 in the Col-0 wildtype background as well as on plants expressing mCit-ROP2 in the *gef3-1* mutant background.

Similar to ROP2, we observed sub-compartmentalization of GEF3 within nanodomains at the RHID using VAEM (Fig.2B) and confocal laser scanning microscopy (Fig.S2B). At cell stage -4, mCit-GEF3 nanodomains localized predominantly to the basal (root-meristem-oriented) end of the cell, but, to a lesser extent, were also found distributed across the cell surface. In subsequent developmental stages, the basal localization of GEF3 nanodomains was gradually lost and GEF3 fully polarized at the RHID. This translocation of mCit-GEF3 from the basal cell-cell contact to the RHID during the earliest stages of root hair initiation has been reported previously (Denninger et al., 2019). Similar to ROP2, the area covered by the GEF3 patch reached its approximate final scale in cell -3 and the density of GEF3 nanodomains remained largely constant over the course of RHID formation. The distinct localization of proteins within nanodomains at the RHID shown for ROP2 in Col-0 background, was mostly lost when looking at mCit-ROP2 in the *gef3-1* mutant background. While ROP2 in the *gef3-1* mutant still localized in nanodomains, these clusters were evenly distributed throughout the PM of the trichoblasts observed.

Taken together, our data suggest that while the localization of ROP2 into nanodomains is independent of GEF3, the restriction of these nanodomains to the RHID requires a direct or indirect interaction between ROP2 and GEF3. Even though the density of nanodomains in the RHID remained constant for ROP2 as well as for GEF3, the area of the RHID increased during cell stage -4 and -3. We therefore hypothesized, that an increase in RHID area is mediated by new nanodomains, which polarize at the RHID with a predefined spacing to already present nanodomains.

### Association with the PM is a prerequisite for ROP2 polarization at the RHID

Electrostatic interactions between polybasic regions of proteins and anionic lipids are known to play a role in the correct targeting of these proteins to their subcellular destination (Platre et al., 2018). It is further known, that anionic lipids such as phosphatidylserine and phosphatidyl-4,5-bisphosphate (PI(4,5)P2) in conjunction with proteins containing polybasic stretches can promote nanodomain formation in membranes (van den Bogaart et al., 2011). In plants, PI(4,5)P2 fulfills important functions in vesicle trafficking during tip growth in pollen tubes and root hairs (Kost et al., 1999; Braun et al., 1999; Kusano et al., 2008; Mendrinna and Persson, 2015; Noack and Jaillais, 2020). Recently, Platre and co-workers have shown that this mechanism is also involved in targeting of ROP6 into phosphatidylserine-mediated membrane nanodomains (Platre et al., 2019). Since ROP2 contains a polybasic tail close to its C-terminus (Fig.S2A), we tested whether interactions with anionic lipids might be involved in ROP2 polarization at the RHID. We hypothesized that, if ROP2-lipid interactions were the driving force for ROP2 polarization, a change in the local lipid composition within the RHID would precede ROP2 accumulation.

We therefore analyzed several reporters for anionic lipids with respect to their localization in trichoblasts prior to and shortly after bulging (cell stage -1 and +1, respectively). For the reporters for phosphatidylinositol (4) phosphate (PI4P) and phosphatidylserine (PS) (Simon et al., 2014, 2016) as well as for a reporter for general membrane surface charge (MSC) (Simon et al., 2016), we could not observe a polar localization in cell stages -1 to +1 (Fig.S3). Recently, we found the reporter for PI(4,5)P2 to polarizes at the RHID in cell stage +1 (Denninger et al., 2019). Together, these data indicate that PI(4)P and PS are not specifically enriched at the RHID and PI(4,5)P2 is enriched only upon the onset of bulging, following the targeted recruitment of its kinase PIP5K3 in cell stage -1 (Denninger et al., 2019).

Since lipid reporters are only available for certain lipid species and the localization of the specific lipid species in question may thus remain elusive, we further investigated a mutant of ROP2 that we predict would lose its affinity to anionic lipids. We generated a *rop2* mutant in which 7 lysines of the C-terminal polybasic tail were substituted by 7 alanines (*rop2* 7K-A, Fig.S4A). Similar to a *rop2* mutant where the C-terminal membrane anchor had been deleted (*rop2*ΔC161, Fig.S4A), *rop2* 7K-A showed a loss in polarity, and furthermore, a reduction in membrane attachment (Fig.S4B,C,D). In addition, neither a ROP2 C-terminus including the polybasic region (*rop2*ΔN160), nor an artificial poly-lysine motive attached to the PM via a farnesyl-anchor (8K-Farn, Simon et al. (2016)) showed polar accumulation at the RHID, but both retained PM association (Fig.S4B,C,D). Consequently, the reduction in membrane association after mutating the polybasic tail of ROP2, which is in line with similar results obtained for ROP6 (Platre et al., 2019), prevents us from testing whether anionic lipids are directly involved in targeting ROP2 into nanodomains.

Taken together, the data presented here suggest that the poly-basic tail of ROP2 is involved in targeting ROP2 to the PM, but is not sufficient for polarization at the RHID. These findings do not exclude a role of anionic lipids in ROP2 polarization but favor a hypothesis where lateral sorting is lipid-independent. Nevertheless, anionic lipids clearly are of central importance for ROP2 PM-recruitment and during targeted secretion once growth is initiated. Furthermore, our data can be interpreted as evidence that a non-polar association with the PM is a prerequisite for lateral sorting in the PM, potentially via GEF3.

### ROP2 mobility is reduced inside the RHID

To test the hypothesis that GTPase polarization at the RHID is achieved by lateral sorting of ROP2 within the plasma membrane, we measured ROP2 protein dynamics by particle tracking. Being based on the blinking of mCitrine, we termed this approach Tracking of Blinking Fluorophores (ToBF). To reduce false allocation of two distinct proteins into the same track, due to high protein density, we decided to link two particles across two subsequent time-frames (interval: 60 ms) only when their distance did not exceed half their diameter (determined as d = 0.5 µm), creating an upper limit of measurable track velocities (of 0.3 µm/60 ms = 5µm/sec). Using ToBF, we observed that a portion of the particles identified were visible for only a single time frame (Fig.3B, purple circles), which was in line with our previous observation of the occurrence of fluorophore blinking of mCitrine (Fig.1, Fig.S1). Analyzing the dynamics of particles that were traceable for longer than two time points, we found that tracks within the RHID were in general slower compared to tracks outside the RHID (Fig.3C).

**Fig. 3.**
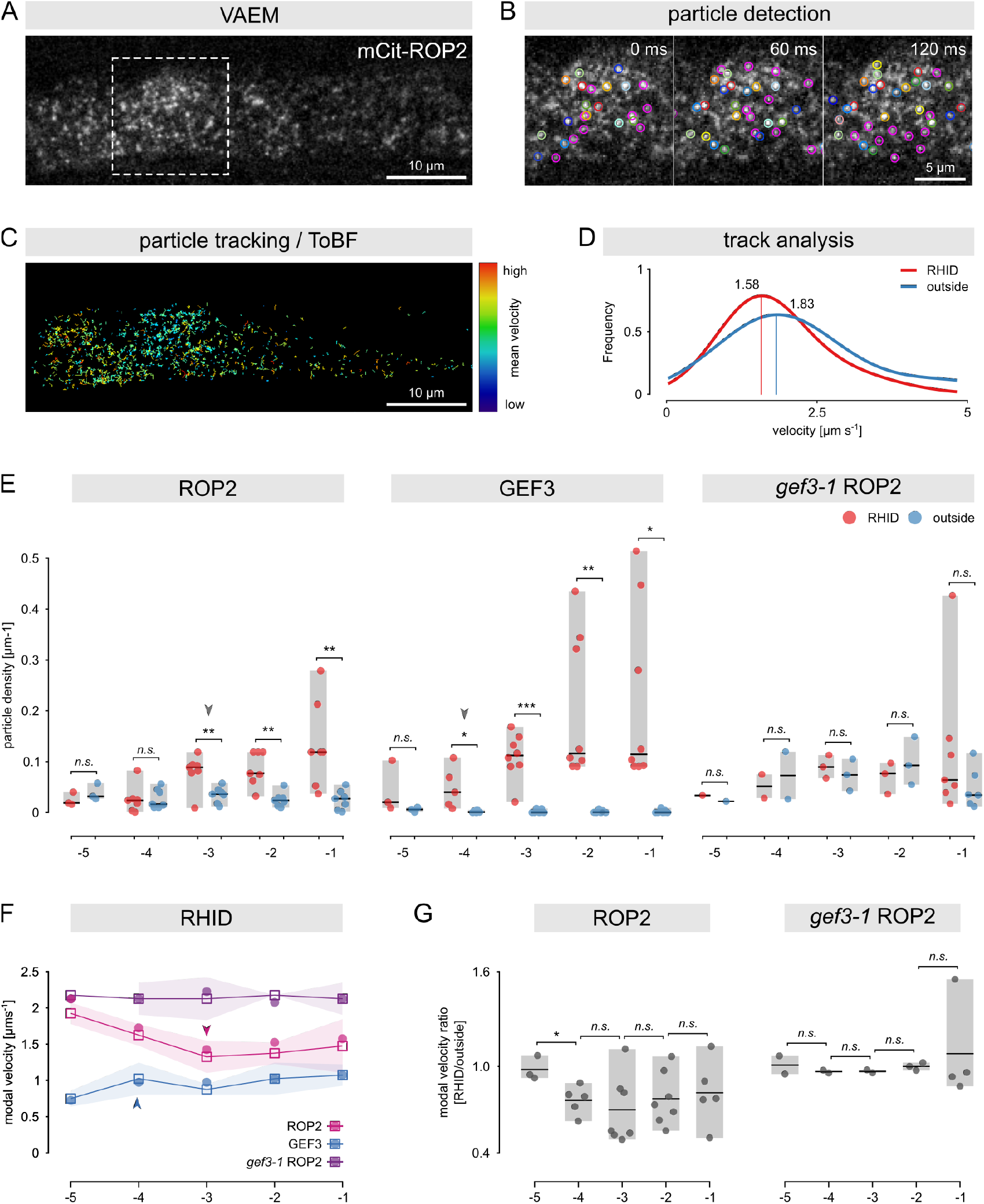
GEF3-dependent reduction in protein mobility precedes ROP2 polarization at the RHID. (A) VAEM-micrograph of a trichoblast of stage -1, expressing mCit-ROP2; the area in the dashed square is enlarged in panel B; scale bar represents 10 µm. (B) enlarged area of a VAEM time-lapse stack indicated by the dashed square in panel A, particles detected with the Laplacian of Gaussian (LoG) detector are indicated by circles. Note: circles of the same color (except purple) were identified in several, subsequent frames of the time lapse stack; purple circles were identified only in one frame; scale bar represents 5 µm (C) result of ToBF (tracking of blinking fluorophores) detected in the in panel A shown VAEM time-lapse stack; tracks are color-coded by their mean velocity; scale bar represents 10 µm (D) normalized distribution of mCit-ROP2 particle velocities, inside (red) and outside (blue) the RHID in a -1 cell. Data was fitted by multi-gaussian fitting; local maxima (“modal velocities”) are depicted by lines and indicated by numbers. (E) particle density determined by particle detection (in) and outside (out) the RHID for ROP2, GEF3 and ROP2 in the *gef3-1* mutant background. Center lines represent median values; gray boxes show the data range. Arrows indicate the first cell stage in which the protein was found to be polar; (F) modal velocities for ROP2, GEF3 and ROP2 in the *gef3-1* mutant background in the RHID of trichoblast cell stages -5 to -1. Arrows indicate the first cell stage in which the protein was found to be polar; circles represent the modal velocity of the pooled data; squares represent the median of modal velocities of individual experiments; shadow represents the MAD (median absolute deviation). (G) Ratio of modal velocities of ROP2 and *gef3-1* ROP2, calculated per individual cell by dividing the modal velocity in the RHID by the modal velocity outside the RHID. Gray boxes represent the data range; center line represents the mean. Asterisks indicate statistically significant differences; p-value determined by Student’s t-test: n.s. = p-value >0.05; * = p-value < 0.05; ** = p-value < 0.01; *** = p-value < 0.001.

To quantify this difference in mobility, we generated histograms of all measured track velocities within a single cell, inside and outside the RHID, and normalized the sample distribution, for better comparison between replicates, to the velocity that occurred with the highest frequency. Data from different biological replicates were integrated by applying multi-gaussian fitting to the collectivity of all histograms. The local maximum of the fitted curve, i.e. the mode of the velocity distribution (or: modal velocity), indicates the velocity with which the majority of particles moved within the PM, and therefore serves as a single-value measure for over-all protein mobility that can be used for comparison between different conditions and/or proteins (Fig.3D, Fig.S5).

We determined mCit-ROP2 protein mobility by Fluorescence Recovery After Photobleaching (FRAP) and found that the mobility of ROP2 inside the RHID was significant lower compared to outside the RHID (median t½ for ROP2 in the RHID 40.27 sec, for ROP2 outside 5.75 sec; p-value 0.01), whereas the immobile fraction at the RHID was slightly higher, but was not statistically significant different (10.80% for ROP2 at the RHID and 0% for ROP2 outside the RHID, p-value 0.97) (Fig.S6A-B). In line with these results, ToBF revealed that the modal velocities of mCit-ROP2 molecules inside the RHID was lower (1.58 µm s^−1^) compared to outside the RHID (1.83 µm s^−1^) (Fig.3D).

Together, the data presented here indicates a local immobilization of ROP2, specifically at the RHID.

### Immobilization of ROP2 precedes its polarization

To investigate to which extent this reduction in mobility of mCit-ROP2 molecules at the RHID might correlate with RHID development, we determined the earliest cell stage in which ROP2 polarizes at the RHID using VAEM time-lapse stacks. The local concentration of FPs (representing the concentration of the protein of interest, in this case ROP2) is positively correlated with FP blinking, corresponding to the presence of an FP in the “On” state. Therefore, we chose the density of particles – which is equivalent to the density of blink events – as a measure for relative local protein abundance. For the analysis of polarity, a protein was then deemed polar when the density of particles in the RHID was significantly higher than the density of particles in the PM outside the RHID.

The earliest cell stage in which mCit-ROP2 polarization occurred at the RHID was stage -3 (Fig.3E). During later developmental stages the density of particles at the RHID did not increase further in a statistically significant manner (p-values for particle density at the RHID: -5 vs -4, 0,47; -4 vs -3, 0.003; -3 vs -2, 0,38; -2 vs -1, 0,1) but remained at a higher level relative to outside the RHID. Recently, we were able to report that GEF3 is necessary and sufficient to polarize ROP2 and precedes ROP2 polarization at the RHID (Denninger et al., 2019). In line with this finding, mCit-GEF3 polarization at the RHID, as quantified by particle density, was detected one cell stage earlier than mCit-ROP2, namely at cell stage -4 (Fig.3E). In the *gef3-1* mutant however, mCit-ROP2 did not show a polar accumulation at the RHID (Fig.3E).

With this information, we proceeded to analyzing the protein mobility over the course of root hair development. We fitted velocity histograms individually for each biological replicate, which allowed us to compare median values of modal velocities (represented as squares in Fig.3F), inside the RHID and for each cell stage. Furthermore, we calculated the ratio of modal velocities between inside and outside the RHID for each cell individually, allowing us to perform statistical analyses and compare different plant cells and cell stages with each other (Fig.3G). A ratio of 1 would mean that proteins inside and outside the RHID moved with the same velocity. Consequently, a ratio below 1 would indicate a reduction in velocity at the RHID.

For mCit-GEF3, no particle could be detected outside the RHID, thus no ratio of modal velocities was calculated. In comparison to mCit-ROP2, the modal velocity of mCit-GEF3 was lower (ranging from 0.7 µm s^−1^ in cell -5 to 1 µm s^−1^ in cell -1) and remained constant during root hair development (Fig.3F). For mCit-ROP2, we found a reduction of modal velocity at the RHID at cell stage -4 (Fig.3F,G), preceding its polarization at cell stage -3 (indicated by the arrow head in Fig.3F). During subsequent root hair development ROP2 mobility inside the RHID increased slightly while its modal velocity ratio remained constant, below 1 and statistically unchanged (Fig.3G). Loss of GEF3 led to an increase of mCit-ROP2 mobility in general (Fig.3F) and a loss of differential immobilization at the RHID (Fig.3G, *gef3-1* ROP2). These findings were consistent with FRAP measurements performed in cell stage -1, which showed that the protein mobility of GEF3 is lower compared to ROP2 (median t½ for ROP2 in the RHID 40.27 sec, for GEF3 104.87 sec; p-value 0.01) and that the reduction in ROP2 protein mobility inside the RHID is dependent on the presence of GEF3 (Fig.S6).

Taken together, we show evidence, that the immobilization of ROP2 in nanodomains at the RHID is dependent on the immobile protein GEF3, whose polarization not only precedes but represents a precondition for ROP2 polarization. Together these findings lead us to conclude that GEF3 facilitates ROP2 polarization by reducing lateral mobility of ROP2 in the PM of the RHID.

### Differential activation of ROP2 required for polarization, but not for nanodomain association

GTPases, like ROP2, act as molecular switches that can be present in an active (GTP-bound) and an inactive (GDP-bound) state. To investigate whether the activity state of ROP2 plays a role in its polarization at the RHID, we determined the polarity index of two ROP2 activity state mutants (*rop2*CA = constitutive active, GTP-locked; *rop2*DN = dominant negative, GDP-locked) in trichoblasts of cell stage -1. We found that neither mCit-*rop2*CA nor mCit-*rop2*DN showed predominant accumulation at the RHID, but that both had polarity indices similar to the unpolar controls GFP-LTI6B (Cutler et al., 2000) and freely soluble, cytosolic mCitrine (Fig.4A,B).

**Fig. 4.**
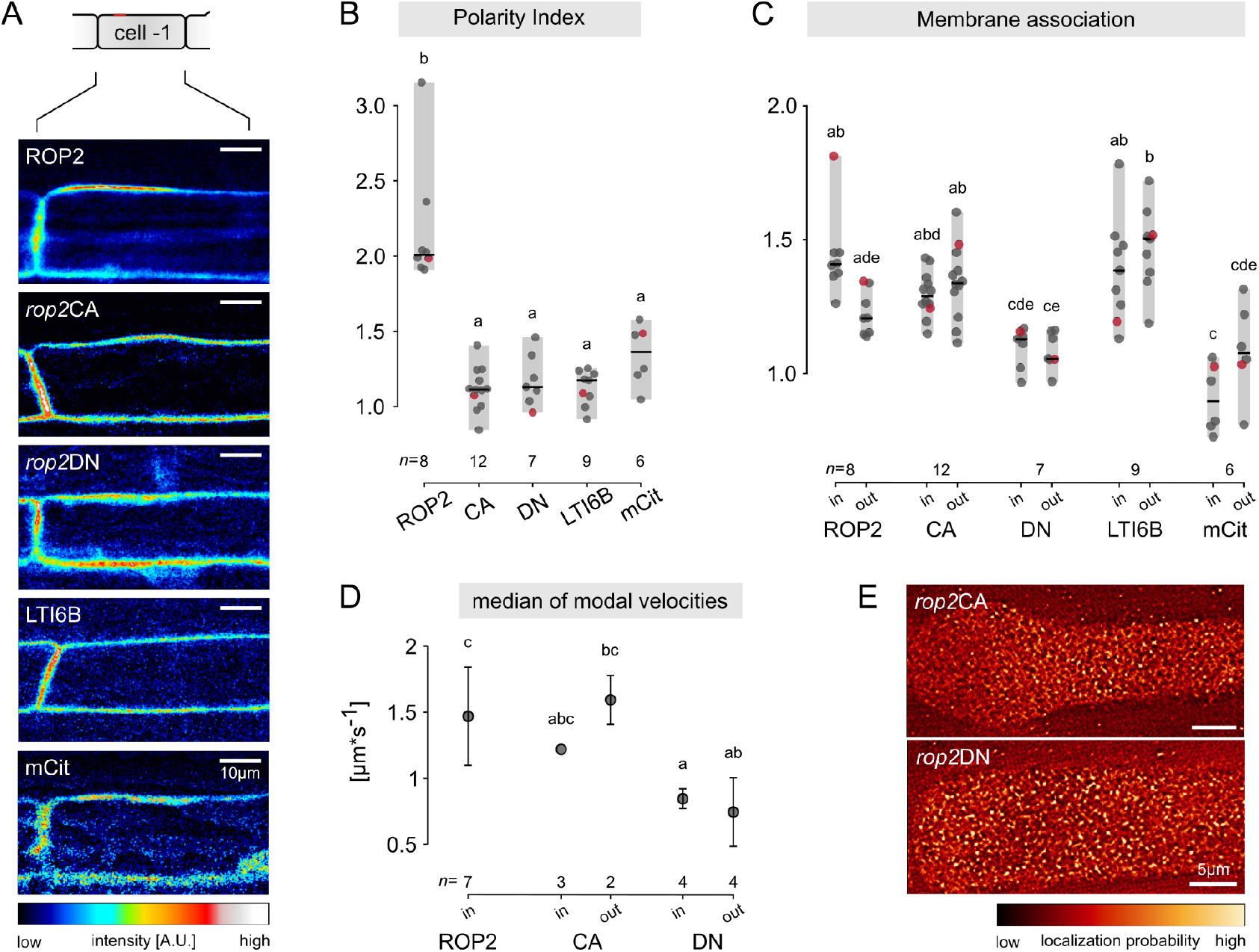
Differential activation is necessary for ROP2 polarity establishment. (A) Micrographs of trichoblasts of cell stage -1 expressing mCit tagged ROP2, mCit-*rop*CA (CA), mCit-*rop*DN (DN) as well as GFP-LTI6B and cytosolic mCitrine (mCit). Scale bars represent 10 µm; the root tip is located to the left of the images. (B) Polarity index and (C) association with the plasma membrane inside (in) and outside (out) of the RHID, of the plant lines depicted in panel A. Centre lines represent median values, gray boxes represent the data range, n indicates the number of cells measured and letters represent the result of an ANOVA-Tukey test (significance value = 0.01; same letters indicate no significant difference). Red data points indicate data derived from the individuals shown in panel A. (D) Quantification of the modal velocities in cells of stage -1 of mCit tagged ROP2, mCit-*rop*CA (CA) and mCit-*rop*DN (DN) inside (in) and outside (out) of the RHID. Circles represent the median of the modal velocities; error bars represent the MAD (median absolute deviation); letters represent the result of an ANOVA-Tukey test (significance value = 0.01; same letters indicate no significant difference). (E) SRRF reconstructions of -1 trichoblasts expressing mCit tagged *rop*CA and *rop*DN; the root tip is located to the left of the images; scale bar represents 5 µm.

Since current models suggest that a cycling between the two activity states would be accompanied by a shift in subcellular localization from the cytosol to the PM (Kawano et al., 2014), we measured the degree of membrane association of both activity state mutants of ROP2 (Fig.4C). In line with the current model, we found the degree of membrane association of mCit-*rop2*CA to be comparable to that of the integral membrane protein LTI6B used as reference. In contrast, the association of mCit-*rop2*DN with the PM resembled the degree of membrane association of the cytosolic control mCitrine (Fig.4C), indicating a partial reduction in membrane association for *rop2*DN. Interestingly, the membrane association of wild type (wt) ROP2 (which represents a mix of ROP2 molecules in their GTP-bound and GDP-bound state) outside the RHID was reduced by 14% compared to inside the RHID. We conclude that the dynamic cycling of ROP2 between both activity states is required to polarize the protein at the RHID. To investigate the impact of the activity status on the local immobilization of ROP2 at the RHID, we performed mobility measurements of both activity state mutants. FRAP experiments at cell stage -1 revealed a reduced mobility of mCit-*rop2*CA compared to wt ROP2 with a larger immobile fraction (10.80% for ROP2 and 44,20% for *rop2*CA, p-value 0.03) (Fig.S7A,C). mCit-*rop2*DN, on the other hand exhibited higher mobility than wt ROP2, but also an increased immobile fraction (48.40% for *rop2*DN, p-value for comparison with ROP2 0.02) compared to wt ROP2 (Fig.S7B,C). Performing ToBF on mCit-*rop2*CA at cell stage -1 confirmed its reduced mobility compared to wt ROP2. Interestingly, we were able to detect and track mCit-*rop2*DN using ToBF and found that its modal velocity was reduced to a greater extend as mCit-*rop2*CA (Fig.4D), which was in line with the relatively high immobile fraction measured in FRAP and indicates a residual localization of *rop2*DN at the PM, that is not reflected in the measurement of the membrane association (Fig.S7B,C).

SRRF reconstructions revealed the presence of mCit-*rop2*CA in distinct nanodomains, which however, in line with the polarity index measurements, did not show predominant accumulation at the RHID but were found evenly distributed across the cell surface (Fig.4E). mCit-*rop2*DN similarly localized into stable nanodomains across the cell surface (Fig.4E), which further indicates the presence of a subpopulation of *rop2*DN molecules residing at the PM.

Taken together, the data presented here shows that differential activation is required for ROP2 polarization at the RHID. The association of ROP2 with nanodomains is, however, independent of its nucleotide state.

## Discussion

Targeted recruitment of the underlying growth machinery to the site of root hair emergence, the root hair initiation domain (RHID), is an essential step in root hair development. The small Rho-type GTPase ROP2 is a central player and early determinant in root hair development (Jones et al., 2002; Molendijk et al., 2001). Spatio-temporal control of its sub-cellular localization to the RHID is key to ensure locally restricted outgrowth of a root hair and to the robustness of hair positioning. Even though much is known about the role of ROPs in root hair development, we still do not fully understand by which mechanism sufficient ROP molecules are delivered to the root hair during growth and how initial accumulation of ROPs at the RHID is facilitated. Two mechanisms seem conceivable: Targeted secretion to the RHID or insertion of ROPs into the PM followed by lateral sorting into the RHID.

In this study, we aimed to investigate the mechanism underlying initial ROP2 polarization at the RHID. To this end we employed Variable Angle Epifluorescence Microscopy (VAEM) and took advantage of fluorophore blinking of mCitrine. We performed time-integrated localization of single molecules and mobility analysis using particle tracking, which we termed Tracking of Blinking Fluorophores (ToBF). In general, the quality of single molecule tracking largely depends on fluorophore density: The lower the fluorophore density, the easier it is to follow a single molecule and thus the more precise the measured velocity reflects actual protein mobility. In methods using photoactivatable fluorescent proteins (FPs), e.g. sptPALM (Manley et al., 2008; Bayle et al., 2021), the density of fluorophores can be tuned by adjusting the power of the activating laser and it is possible to obtain relatively long tracks. In contrast, FP density in ToBF cannot be tuned, is generally higher and varies over the course of a time-lapse measurement, as well as between different samples. To still be able to track single molecules at a high particle density, we have allowed for a maximum travelling distance of 0.3 µm between single time frames, thus creating an upper limit of measurable velocities (5 µm s^−1^). Moreover, tracking of blinking FPs is only possible as long as the FP is in the excitable and emitting “on” state, which is a stochastic process and generally leads to shorter tracks. While these limitations results in a lower tracking resolution in comparison to sptPALM, our ToBF analysis allows for the qualitative measurement of mobility on the scale of the whole protein population, utilizing a standard fluorophore that is already commonly used in many labs and existing reporter lines.

In addition to protein mobility, fluorophore blinking allows to perform time-integrated single molecule localization, which has led to techniques such as dSTORM (Heilemann et al., 2008) or GSDIM (Fölling et al., 2008). Here, we subjected our raw VAEM data of blinking fluorophores to the Super Resolution Radial Fluctuation (SRRF) algorithm (Gustafsson et al., 2016), which is suitable for the deconvolution of timelapse datasets based on fluorophore intensity fluctuations. SRRF performs a time-integration of single molecule localization events, where high positional stability (equal to localization probability) is reflected by high pixel values, providing a visual representation of protein mobility. Since SRRF can be applied on images of unfixed, living tissue, the resulting increase in resolution is limited by protein movement and the temporal resolution that can be achieved. The combination of VAEM with SRRF allows for faster acquisition of super-resolved images compared to other super-resolution techniques: A single SRRF image is computed via the integration of 100 frames of a VAEM time-lapse stack, which, at an exposure time of 60 msec per frame, represents a total acquisition time of 6 seconds. However, the stochastic nature of fluorophore blinking also limits the possibility of colocalization studies on mobile proteins, as statistical analyses depend on largely simultaneous emission of spatially correlated fluorophores. However, the compatibility with standard FPs and live imaging enables the visualization of dynamic, cellular processes, e.g. during cell growth, over time. In summary, the combination of VAEM with ToBF and SRRF allows to investigate cellular processes at increased spatial resolution without sacrificing temporal resolution and can be used to analyze the dynamic behavior of a protein population utilizing a commonly used standard FP.

Applying SRRF on trichoblasts expressing mCit-tagged ROP2, we observed the localization of ROP2 into nanodomains localizing at the RHID after its polarization in cell stage -3 (Fig.2 A,B). In line with previous findings (Denninger et al., 2019), we found the polarization of ROP2 nanodomains at the RHID to be dependent on GEF3, while their association with nanodomains *per se* was not (Fig.2C). At the same time, by applying ToBF, we observed a GEF3-dependent reduction in lateral mobility for ROP2 specifically in the PM of the RHID, one cell stage prior to its polarization (Fig.3F). Together, these data suggest a step-wise process for ROP2 polarization at the RHID: First, ROP2 associates with the PM, which involves interaction with anionic lipids, second, ROP2 is sequestered into membrane nanodomains, and third, ROP2 nanodomains are then laterally sorted into the RHID. Such a three-step mechanism would allow for preloading of ROP2 within the PM at a sub-critical concentration before root hair initiation, enabling for the rapid formation of root hairs in response to signaling or environmental cues.

The finding that GTPase polarization is initiated via lateral sorting within the PM addresses a long-standing question in cell morphogenesis: How are polar sites of growth established? Current models, e.g. for PIN polarity, suggest that polar protein accumulation can be maintained by polar secretion, selective endocytosis and a reduction of lateral mobility through association with nanodomains (Kleine-Vehn et al., 2011). In 2019, Gendre et al. provided evidence that polar secretion of ROP2 is required for the maintenance of the ROP2 patch and for successful root hair outgrowth. The authors demonstrated that the TGN-localized proteins YIP4a and YIP4b are involved in the activation as well as the accumulation of ROPs at the PM of root hairs (Gendre et al., 2019). Even though the amount of ROP2 at the RHID was reduced in the *yip4ayip4b* double mutant and root hair growth was mostly abolished, ROP2 was still present at the RHID in such mutant plants. This indicates that while targeted recruitment may be involved in the maintenance of ROP2 polarization, it is unlikely to be the mechanism responsible for the initial accumulation of ROP2 at the RHID.

Cytoskeletal organization and anionic lipids are both known drivers for targeted secretion (Kost et al., 1999; Braun et al., 1999; Kusano et al., 2008; Mendrinna and Persson, 2015; Noack and Jaillais, 2020) and thus were also promising candidates to guide ROP2 polarization. In root hair development, we have, however, no evidence that organizers and nucleators of the actin cytoskeleton are accumulating at the RHID prior to ROP2 (Denninger et al., 2019). Recent publications have highlighted interactions between small GTPases and the anionic lipids phosphatidylserine (PS) and phosphatidylinositol 4,5bisphosphate (PI4,5P2), both of which are also present in nanodomains (Platre et al., 2019; Fratini et al., 2021). In addition, Platre and co-workers have demonstrated that the stability of ROP6 within nanodomains in the PM of root epidermal cells is dependent on the presence of PS, while Fratini and co-workers found no interdependence of NtRac5 with PI(4,5)P2. With the data presented in the present study, we cannot fully exclude the possibility that the localization of ROP2 into nanodomains similarly depends on electrostatic interactions with anionic lipids. However, we could demonstrate that: (**i**) reporters for the anionic lipids did not polarize at the RHID prior to bulging (Fig.S4), suggesting that these anionic lipids do not serve as landmarks for RHID formation; (**ii**) a ROP2 mutant, which was impaired in its ability to interact with anionic lipids exhibited a reduced PM-association in general (Fig.S4), indicating that membrane association (supported by the interaction with anionic lipids) is a prerequisite for polarization. In addition, the polarization of ROP2 precedes the polar accumulation of PIP5K3, as well as of PI(4,5)P2 (Denninger et al., 2019), further indicating that anionic lipids play a role in root hair growth, downstream of RHID set-up.

We further investigated whether ROP activation could dictate the GTPase polarization. In root hairs, polar ROP2 localization is indeed coupled to its nucleotide-binding state, however, rather in an indirect manner, as the following observations will demonstrate. Lack of the ROP inhibiting GDI SCN1 (Carol et al., 2005), as well as overexpression of the RopGEF GEF3 (Denninger et al., 2019), both resulted in multiple polarization events, leading to multiple bulges or branched hairs. One could therefore expect that enhanced ROP GTPase activation is the cause for the multiple polarization events. However, neither *rop*CA (GTP-locked), nor *rop*DN (GDP-locked) exhibited any detectable polar domain formation, suggesting that not the nucleotide state itself, but rather the ability to dynamically switch between both states is necessary for ROP2 polarization. Our velocity analysis revealed, that in comparison to wild type ROP2, mCit-*rop*CA exhibited reduced mobility and was associated with nanodomains evenly distributed in the PM in cell stage -1 (Fig.4D,E). To our surprise, we were able to also detect mCit-*rop*DN both in the cytosol and in nanodomains at the PM and found the protein mobility at the PM to be even lower than the mobility of wt ROP2 and mCit-*rop*CA nanodomains. *rop*DN is unable to release GDP from its binding pocket and is thus, not activatable. It seems conceivable, that this results in constant binding to the inhibiting, cytosolic GDI or its PM-bound activators and thus causes their sequestration, leading to its eponymous, dominant negative phenotype (Glotzer and Hyman, 1995; Berken and Wittinghofer, 2008). This gives rise to an interesting hypothesis in which GEF3 and SCN1 have to dynamically compete for an interaction with the GTPase, where GEF3 serves as polar landmark at the PM while cytosolic SCN1 counteracts an overloading of the RHID with ROPs. On the one hand, *rop2*DN may be mostly unable to dissociate from SCN1, thereby unable to follow GEF3 localization, while a subpopulation of *rop*DN being trapped in static physical interactions at the PM. On the other hand, *rop*CA localized to immobile PM nanodomains, and over all showed a reduced mobility compared to wt ROP2, but an increased mobility compared to *rop*DN (determined by ToBF, Fig.4D).

Although not all questions regarding the robust polarization of ROP GTPases during root hair morphogenesis can be answered at this point, our study provides new insights in the protein dynamics of ROP2 and the involved classes of potential drivers and interaction partners. In summary, our findings support a model in which ROPs are first associated with the PM through the interaction with anionic lipids and independent on the bound nucleotide. At the PM, GTPases are locally immobilized in nanodomains and accumulate at the RHID in a GEF3-dependent manner, leading to the establishment of a new polarity axis. In a feed-forward loop, downstream effectors of ROP signaling are recruited, promoting cytoskeletal reorganization, lipid modification and the establishment of targeted secretion that, further on, stabilizes cell polarity and sustains the elongation of the growing root hair. At this point, we can only speculate whether a similar series of events is at play during morphogenetic programs of other eukaryotes with often way more sophisticated cellular architectures. In baker’s yeast, mechanisms involving spontaneous accumulation of the small GTPase Cdc42 that lead to local cytoskeletal reorganization and cell polarization have been identified (Wedlich-Soldner and Li, 2003), highlighting the need for landmarks guiding the formation of polarity axes (Chiou et al., 2017). Also, the morphological robustness of root trichoblasts likely involves preventing a spontaneous polarization at random sites at the PM. As mentioned before, root hairs emerge consistently at the basal (root-meristem-oriented) end of the cell with a consistent diameter, despite the absence of any apparent physical boundaries. Fischer and colleagues identified a dependence of the positioning of the polar domain on auxin signaling pathways (Fischer et al., 2006) giving rise to a mathematical model that suggested the presence of an intracellular auxin gradient that drives ROP polarization through a Turing-like, reaction-diffusion mechanism (Payne and Grierson, 2009). Indeed, in epidermal cells of the root elongation zone, ROP6 was shown to be immobilized upon auxin treatment in PS-dependent nanodomains, which were required for inhibition of cell elongation and consequential gravitropic root bending (Platre et al., 2019). In a recent study on the maintenance of ROP polarization during root hair growth, ARMADILLO REPEAT ONLY (ARO) proteins were found to stabilize the interaction between ROP1 and the ROP1 enhancer GAP (RENGAP) REN1 in PM nanodomains, thus confining ROP signaling to the polar growth site (Kulich et al., 2020). Although nanodomain association had not yet been considered in the Turing-like model by Payne and Grierson 2009, our findings are generally in line with a reaction-diffusion mechanism as we demonstrate that a mobility-regulating nanodomain-association plays a role already during the initial steps of ROP polarization, driving the lateral sorting of GTPases and thereby leading to the formation of the RHID.

Together, the various recent publications on nanodomains in the plant PM highlight their roles in organizing membranes into functionally distinct sub-compartments on the micro- and nanoscale. Likely, a great diversity of nanodomains exists, whose composition, structure and dynamics remain to be explored (Gronnier et al., 2018). Novel microscopic techniques with super-resolving capacities, not only regarding the spatial but also the temporal domain, will be needed to elucidate the molecular dynamics that organize biochemical functions and drive signaling and cellular morphogenesis.

## AUTHOR CONTRIBUTIONS

VAFF and GG conceived the project; VAFF, PD and GG planned the experiments; VAFF and PD performed experiments; YJ provided the lipid reporter lines and suggested experiments; UE provided advice on the imaging, data analysis and interpretation; VAFF, MZ and GG analyzed data; VAFF prepared figures; VAFF and GG wrote the manuscript with input from all co-authors; and all authors read and approved the final version of the manuscript.

## ACKNOWLEDGEMENTS

We thank Karin Schumacher (Heidelberg) for mentorship, advice and access to the Leica SP5, the Nikon Imaging Center at the Heidelberg University for providing the TIRF microscope and technical support, Jan Felix Evers for providing the spinning disc microscope, technical support and advice on image acquisition and analyses and all past and current members of the Grossmann lab for their support and lively discussions. This work was supported by CellNetworks research group funds, research grants by the Deutsche Forschungsgemeinschaft (DFG; GR4559-3-1, GR4559-5-1), funds by the Center of Excellence on Plant Sciences (CEPLAS) and a Heisenberg Professorship to G.G. (GR4559-4-1).

## Materials and Methods

### Plant growth and handling

Plants were either grown on soil or on ½MS-plates (Murashige and Skoog Minimal Organic Powder Medium (Serva), 0.1% MES, pH 5.7, 0.8% plant agar (Duchefa)) under long day conditions (16h light/8h dark) at 21°C.

Induction of the estradiol inducible (EstInd) promoter (described in detail in Denninger et al. (2019) was performed using small stripes of cellulose tissue, imbibed in ½MS containing 20 µM ß-Estradiol (20 mM stock in absolute ethanol) and 0.01% Silwet L77. To investigate the subcellular localization of proteins expressed under the control of an EstInd promoter, induction was performed 3-6h prior to sample preparation and for a duration of 30min. In order to get sufficient fluorescence signal, the time between induction and microscopy varied between plant lines, but was kept as short as possible to minimize overexpression artefacts.

### Plant material

In this study, *Arabidopsis thaliana* of the ecotype Columbia (Col-0) was used as wildtype. Transgenic lines expressing pROP2::mCitrine-ROP2(ORF), EstInd::mCitrine-ROP2(CDS), pGEF3::mCitrine-GEF3 and pROP2::mCitrine-ROP2 in the *gef3-1* background were obtained from Denninger et al. (2019). Lipid reporters (PI(4,5)P2 reporter – P15Y, PI(4)P-reporter (pUbi10::mCitrine-P4M^*SiDM*^), PS-reporter (pUbi10::mCitrine-C2^*Lact*^)) as well as 8K-Farn (pUbi10::mCitrine-8K-Farn) and the MSC sensor (pUbi10::mCitrine-KA1^*MARK*1^) (Simon et al., 2014, 2016) were kindly provided by Yon Jaillais (Lyon). P35S::GFP-LTI6B (Cutler et al., 2000) was kindly provided by David Ehrhardt (Carnegie Institution for Science.). Plant lines expressing pUbi10::mCitrine, EstInd::mCitrine-rop2-7K-A, EstInd::mCitrine-rop2ΔC161, EstInd::mCitrine-rop2ΔN160, EstInd::mCitrine-rop2CA and EstInd::mCitrine-rop2DN were generated in this work.

### Molecular cloning

All expression vectors were cloned using the GreenGate system (Lampropoulos et al., 2013). The estradiol inducible promoter, as well as the mCitrine module was described in Denninger et al. (2019).

LifeAct (Riedl et al., 2008) was cloned into the Green-Gate entry vector pGGC000 by amplifying LifeAct together with a C-terminal GDPPVAT-Linker from the Addgene vector # 36201 using the forward primer aacaGGTCTCtGGC-Tatgggcgtggccgacctgat and the reverse primer acaaGGTCT-CaCTGAggtggcgaccggtggatc.

mNeonGreen (Shaner et al., 2013) was cloned into the GreenGate entry vector pGGD000 by amplifying mNeonGreen from the plasmid pGGD044 (kindly provided by Jan Lohmann, Heidelberg) using the forward primer aaaaGGTCTCaTCAGGAGCAGGGGCGGGT-GCCatggtgagcaagggcgag and the reverse primer aaaaG-GTCTCaGCAGttacttgtacagctcgtcca, indroducing an N-terminal SGAGAGA-linker.

The pUbi10::mCitrine plant expression vector was assembled with the following GG-modules: pGGA6 (Ubip), pPD160 (mCit), pPD37 (C-Decoy), pGGD2 (D-Decoy), pPD59 (HSP18.2-T), pGGF1 (Basta) into pGGZ003. For creation of the C-Decoy module (pPD37), the primer pair forward: gtgaagcttGGTCTCaGGCTgtggatcc and reverse: GC-GAgaattcGGTCTCaCTGAggtacca were used for PCR amplification from the pGGD2 vector. The resulting PCR product was cloned into pGGC000.

ROP2 mutation constructs (*rop2* 7K-A, *rop2*ΔC161, *rop2*ΔN160, *rop2*CA and *rop2*DN) were generated using the CDS sequence of ROP2 as a template and were cloned into pGGC000 using the listed primers (Table S1). For *rop2*DN two PCR-fragments were fused and cloned into pGGC000 by adding both digested PCR products into the ligation reaction.

### Confocal laser scanning microscopy and FRAP measurements

For laser scanning microscopy a Leica SPII system was used, equipped with a 63x water immersion objective (N.A. 1.2, Leica), an argon laser and hybrid-detectors. mCitrine tagged proteins were excited at 514 nm and emission was detected between 520 and 550 nm.

High-resolution surface views of trichoblasts were recorded using the resonant scanner. Images were acquired with a 4x zoom, a pinhole of 1 AU, scan speed of 8000 Hz, 256x line average and a 512×512 px scan field. These settings were kept constant between experiments.

For FRAP the “zoom-in”-modus with 30 frames pre-bleach, 20 frames bleach (100% relative laser power, ROI of 10×30 µm), 200 frames post-bleach (each at 0.265 sec per frame) was chosen. Images were acquired with a 3x zoom, a pinhole of 2.5 AU, at 1000 Hz bidirectional scan speed, 2x line averaging and a 256×256 px scan field. The settings were kept constant between experiments, except for the power of the excitation laser, which was adjusted according to the expression strength of the sample. Analysis of FRAP data was performed by measuring the fluorescence intensity within three ROIs in each time-lapse stack: one ROI at the bleached region, one ROI outside of the biological sample (background) and one ROI at a non-bleached region of the plasma membrane. In each ROI, the fluorescent intensities were normalized to the average fluorescent intensity prior to bleaching. The resulting data set was analyzed using the web-based application “EasyFRAP-web” (https://easyfrap.vmnet.upatras.gr) as described in (Koulouras et al., 2018).

### Sample preparation for VAEM-imaging

Comparison of the blinking behavior of different fluorescent proteins was analyzed in transfected *Nicotiana benthamiana* leaves 2 days after infiltration. Agrobacterium-mediated transfection was performed as has been described in Mehlhorn et al. (2018). To remove air from the intracellular space, tobacco leaves were infiltrated with water shortly before sample preparation and imaging. Small pieces were cut out with a scalpel and placed on a microscopy slide, with the abaxial side facing upwards. A drop of water was added on top and a cover slip (thickness 170 ^+^/_−_ 5 µm; No. 1.5H) was gently pressed onto the sample, using a soft piece of rubber.

Imaging of *Arabidopsis thaliana* root cells was performed 5-7 days after germination. For sample preparation whole seedlings were placed on a cover slide with the root residing in a drop of medium (approximately 20 µL). For imaging, the roots were then carefully covered by a 22×22 mm cover slip (thickness 170 ^+^/_−_ 5 µm; No. 1.5H), whereat the cotyledons were not covered to prevent tilting of the cover slip.

### Variable Angle Epifluorescence Microscopy (VAEM) imaging

For VAEM, a Nikon TIRF microscope was used with a single mode laser fiber coupled into the TIRF illuminator on a Nikon eclipse Ti2 stand to focus the laser on the back focal plane of 100x DIC Plan Apo objective (NA 1.45; Nikon). Only trichoblasts close (few µm) to the coverslip–medium interface where used (as detected by the hardware perfect focus system (PFS) of the Ti2). Detection was performed on an EMCCD camera (Andor iXON Ultra, Andor Technology) with 60 ms exposure time. Continuous recording was performed for 1 min. During acquisition, the hardware focus PFS was enabled. Microscope hardware was controlled by NIS-Elements (version 5.1, Nikon).

Excitation of mNeonGreen and mCitrine tagged proteins was performed using a 515 nm laser and a 542/27 bandpass filter. For excitation the 515 line of an Andor Light Combiner (ALC, Andor Belfast) was used. Fluorophores were driven into the dark state by setting the laser power to 40% (corresponding to 10.7 mW, measured at the objective plane) of the 515 nm laser line. We estimate that this corresponds to approximately 100-150 W/cm^2^ (given the illumination area of the laser through the 100x objective).

### SRRF reconstructions and measurement of nanodomain dimensions

Single-molecule localization was performed by applying the SRRF algorithm (Gustafsson et al., 2016) on VAEM time-lapse stacks. The build in “Estimate Drift”-tool was used with a time-averaging of 100 to estimate the drift prior to SRRF. SRRF reconstruction was performed using the following parameters: ring radius of 0.5, radial magnification of 5, axis ring of 8, 100 frames per timepoint and temporal radiality average (TRA). Additionally, intensity weighing was enabled.

Nanodomain density were measured using Fiji (Schindelin et al., 2012). To this end, a region of interest (e.g. the RHID) in the original SRRF image was thresholded (with the build-in threshold presetting “moments”) and used to create a mask. This mask was processed to a binary image and then multiplied with the original SRRF image. In the resulting image (called “SRRFmasked”), individual nanodomains were identified using the “find maxima” function of Fiji. The output type was chosen as “segmented particles” and the noise tolerance was adjusted manually by verifying the selection in the original SRRF image. The resulting image was converted into a binary image and multiplied with “SRRFmasked”, resulting in the separation of the individual nanodomains. The threshold was adjusted (using the presetting “moments”) and the function “analyze particles” was applied, resulting a table containing the spatial information of each nanodomain. The density of nanodomains was obtained by dividing the number of clusters by the total area of the RHID.

### Tracking of Blinking Fluorophores (ToBF)

For particle tracking a rolling ball background subtraction (radius = 5 px) was performed on the original VAEM image stack, followed by three iterations of a gaussian blur filter (sigma = 0.5). The gray value of each pixel of the resulting image was divided by the maximum gray value of the overall image. This image stack was then multiplied by the original, unprocessed time lapse stack and converted to a 16-bit image while retaining the full gray scale range.

Particle tracking was performed using the Fiji-plugin Track-Mate (Tinevez et al., 2017). Particle detection was performed with the Laplacian of Gaussian (LoG) detector, using an estimated blob diameter of 0.5 µm (sub-pixel localization was enabled). To prevent splitting and merging of tracks, the simple linear assignment problem (LAP) tracker was used. The maximum linking, as well as the maximum gap-closing distance were set to 0.3 µm. Gap-closing was prevented by using no frame gap.

To determine the velocity of proteins, the “mean speed” parameter of the particle tracking was used for further analysis. A histogram of velocities (bins: 0.05µm/sec) was generated and subjected to non-linear least-square fitting using the Gauss-Newton algorithm. Fitting was done with the nls() function, while predicted fittings with corresponding local peak maxima’s were plotted with the ggplot2 and ggthemes packages (Wickham, 2016; Arnold, 2019). To test the differences between the RHID and the PM region outside the RHID, the raw data for each region was used to fit the linear model with the lm() function. Further comparison of fitted models for the different regions was performed with the Anova() function from the car package (Fox and Weisberg, 2019). All analyses were done with R Studio version 1.3.1093 (RStudio Team, 2020).

### Polarity-index and membrane association measurements

Live-cell imaging of transgenic Arabidopsis roots was performed using a custom-built spinning disc confocal microscope (as described in Denninger et al., 2019). mCitrine tagged proteins were excited with a 515 nm laser and emission was detected using a 542/27 (central wavelength/ band width in nm) bandpass filter (Semrock). GFP-tagged proteins were excited with 488 nm and emission was detected using a 525/45 band width filter (Semrock).

Polarity indices were measured and calculated as previously described in Denninger et al., 2019. In brief, the polarity index is the quotient of the background subtracted fluorescent intensity in a 3×15 pixel ROI at the plasma membrane (PM) and the background subtracted fluorescent intensity at the PM outside the RHID [(Int_*PM*−*RHID*_-Int_*Bkgd*_)/(Int_*PM*−*out*_- Int_*Bkgd*_)].

The degree of membrane association of fusion proteins was analyzed by measuring the mean fluorescence intensity in a region of 3×15 pixel at the PM. From this, a background value measured outside the biological sample was subtracted. Next, the mean fluorescence intensity in a region directly underneath the PM-region, in the cytosol, was measured. Background subtraction was performed with the same mean fluorescence intensity as for the PM-ROI. Membrane association was determined by dividing the value of the PM-ROI by the value of the cytosol-ROI [(Int_*PM*_ -Int_*Bkgd*_)/(Int_*Cyt*_- Int_*Bkgd*_)].

## Supplemental Material

**Table S1.**
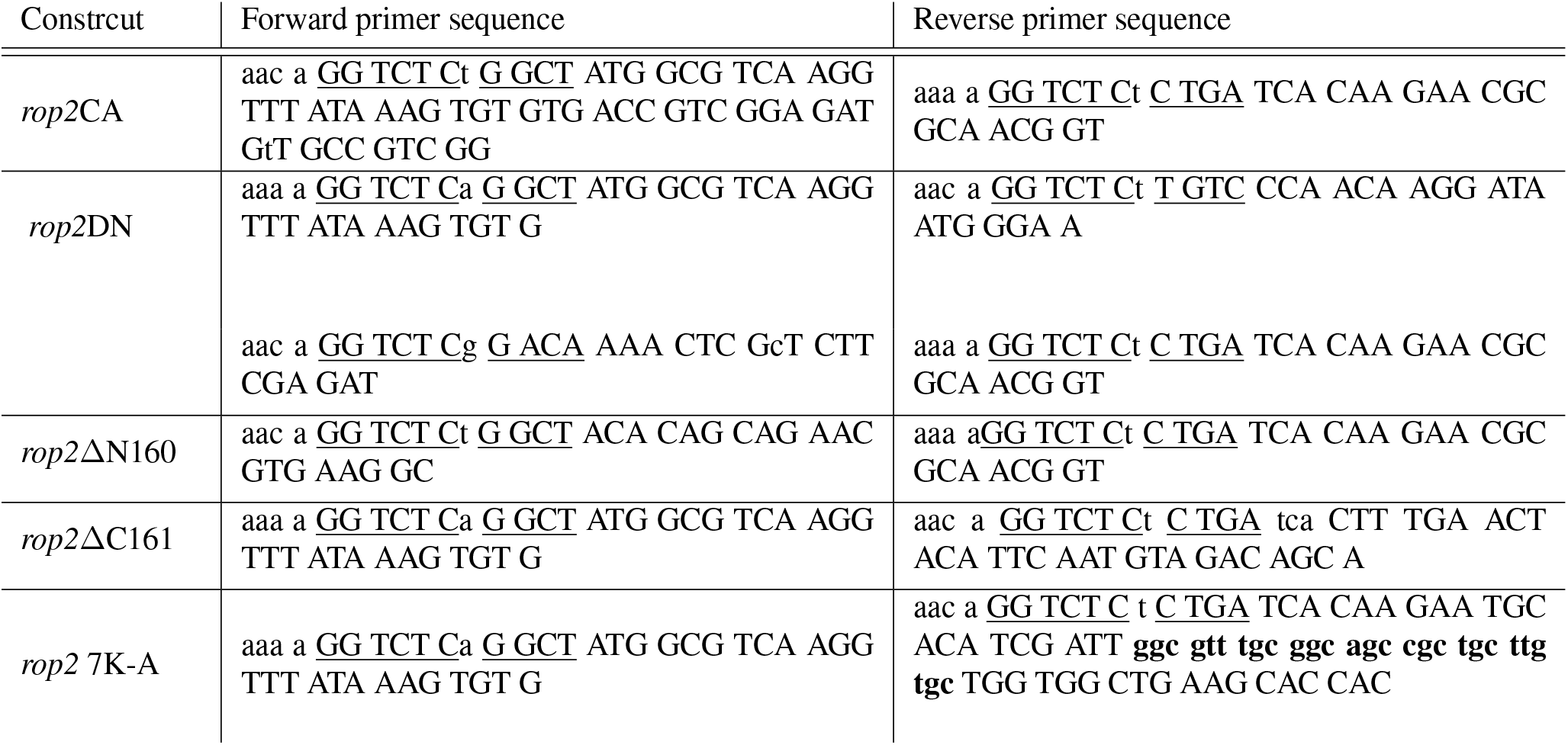
primers used for cloning ROP2 mutant constructs. BsaI/Eco31I recognition and overhang are underlined, mutated triplets are bold and altered bases are in lower case.

**Fig. S1.**
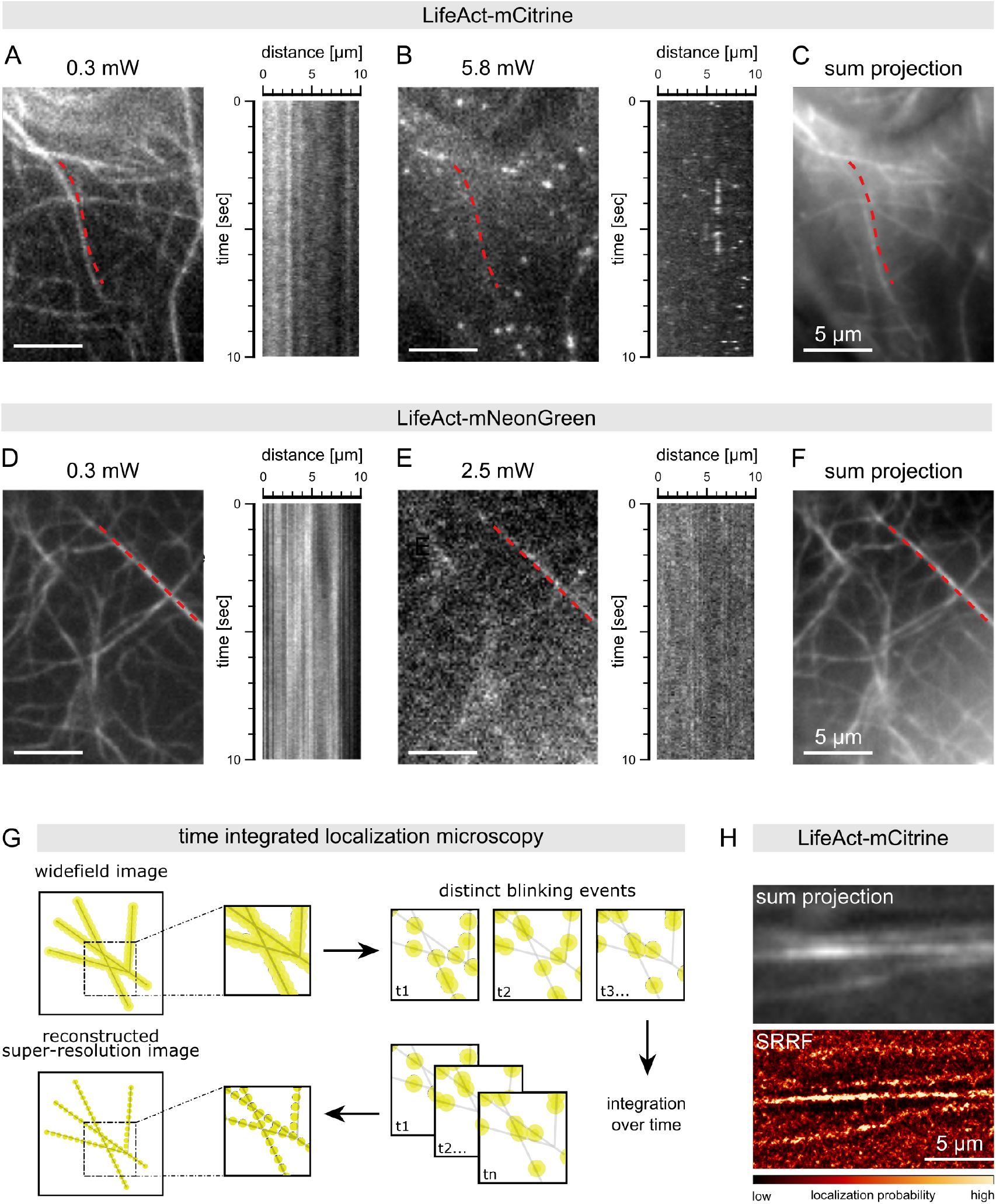
mCitrine exhibits fluorophore blinking, allowing for time-integrated localization microscopy. VAEM-micrographs of the actin probe LifeAct tagged with mCitrine (A-C) and mNeonGreen (D-F). Single images from a time laps stack acquired with the indicated laser powers. Kymographs drawn along the red, dashed lines are depicted next to the corresponding image. Panel (C) and (F) show sum projections of the time laps stacks of (B) and (E), respectively. Scale bars represent 5 µm. (G) schematic representation of the principle of single molecule localization microscopy: the detection and integration of distinct blinking events over time allows for the reconstruction of a super-resolved image (H) Sum intensity projection (upper panel) and a Super-resolution radial fluctuations (SRRF) reconstruction (lower panel) of a time-laps VAEM movie of an Arabidopsis trichoblast stably expressing LifeAct-mCitrine under the control of the *Ubiquitin10* promoter. The scale bar represents 5 µm.

**Fig. S2.**
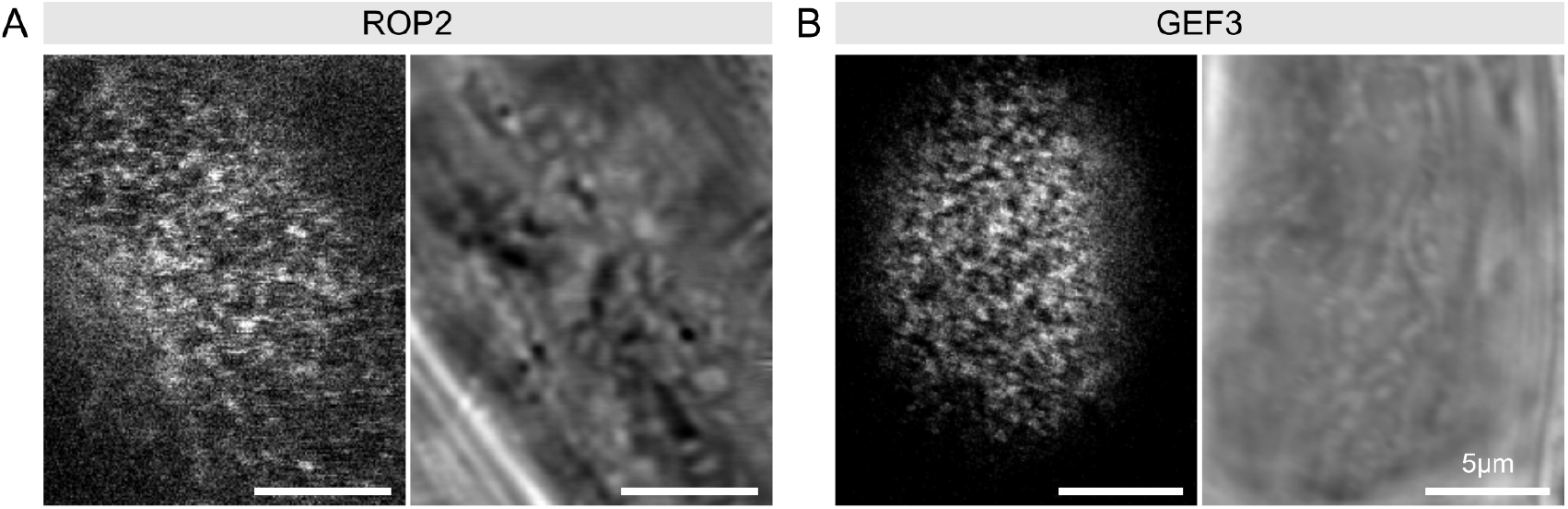
High-resolution laser scanning confocal microscopy cofirms sub-compartmentation of the RHID. Micrographs of mCit-tagged ROP2 (A) and GEF3 (B) in surface-view of trichoblasts of cell stage -1. Scale bar represents 5 µm, the scanning direction was from left to right; the root tip was located to the bottom of the images.

**Fig. S3.**
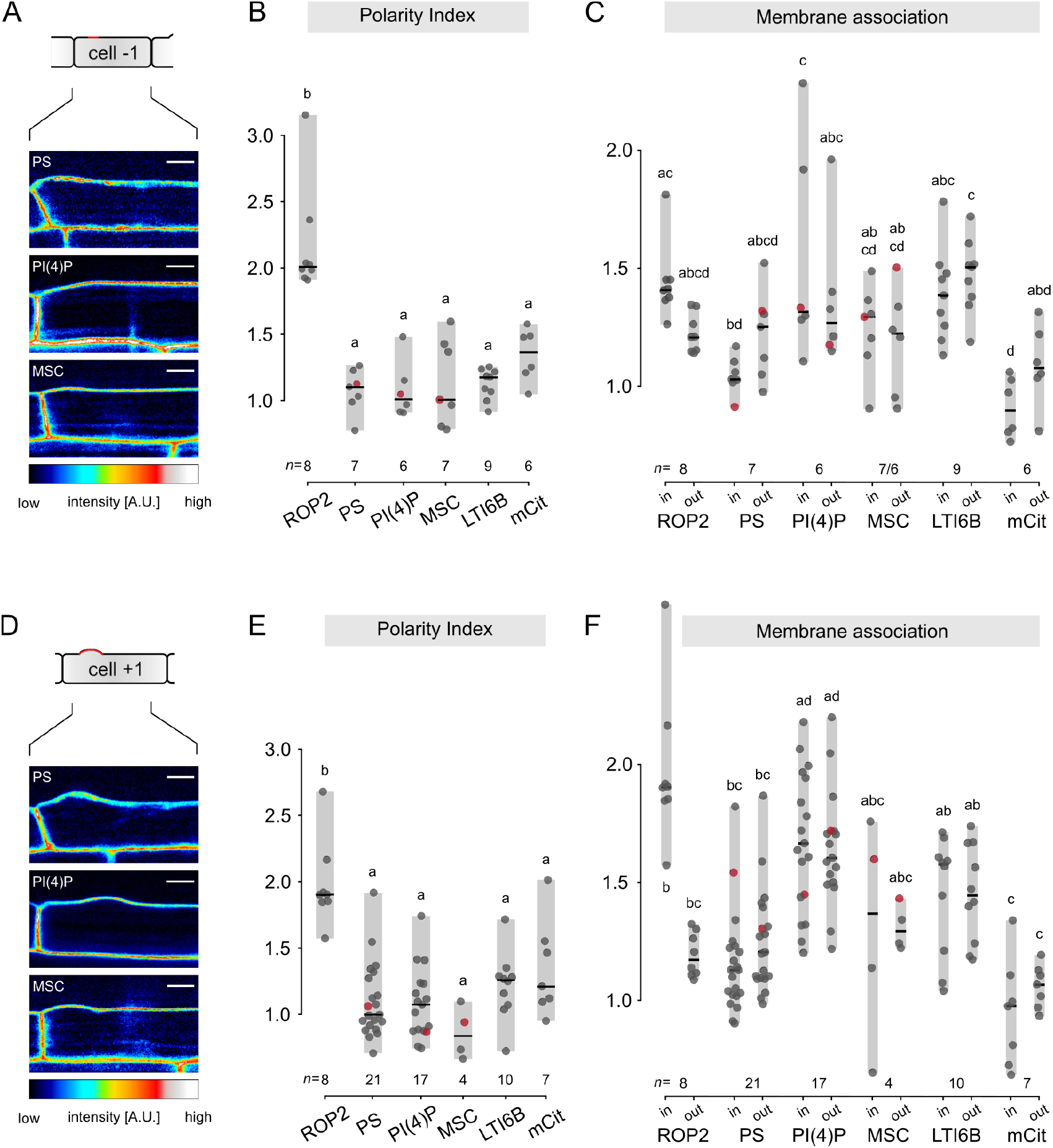
Anionic lipids do not polarize at the RHID prior to bulging. Micrographs of trichoblast cells of stage -1 (A) and of cell stage +1 (D) from plants expressing different markers for anionic lipids – phosphatidylinositol 4-phosphate (PI4P), phosphatidylserine (PS) and mCit-MARK1 (sensor for Membrane surface charge (MSC)=anionic lipids in general). Scale bar represents 10 µm. Quantification of polarity index (B, E) and membrane association (C,F) of the respective markers for anionic lipids in stage -1 (B,C) and in cell stage +1 (E,F). Data for ROP2, mCit and LTI6B are the same as presented in Fig.4, but are shown for comparability. Centre lines represent median values, gray boxes represent the data range, n indicates the number of cells measured and letters represent the result of an ANOVA-Tukey test (significance value = 0.01; same letters indicate no significant difference). Red data points indicate data derived from the individuals shown in panel A and D.

**Fig. S4.**
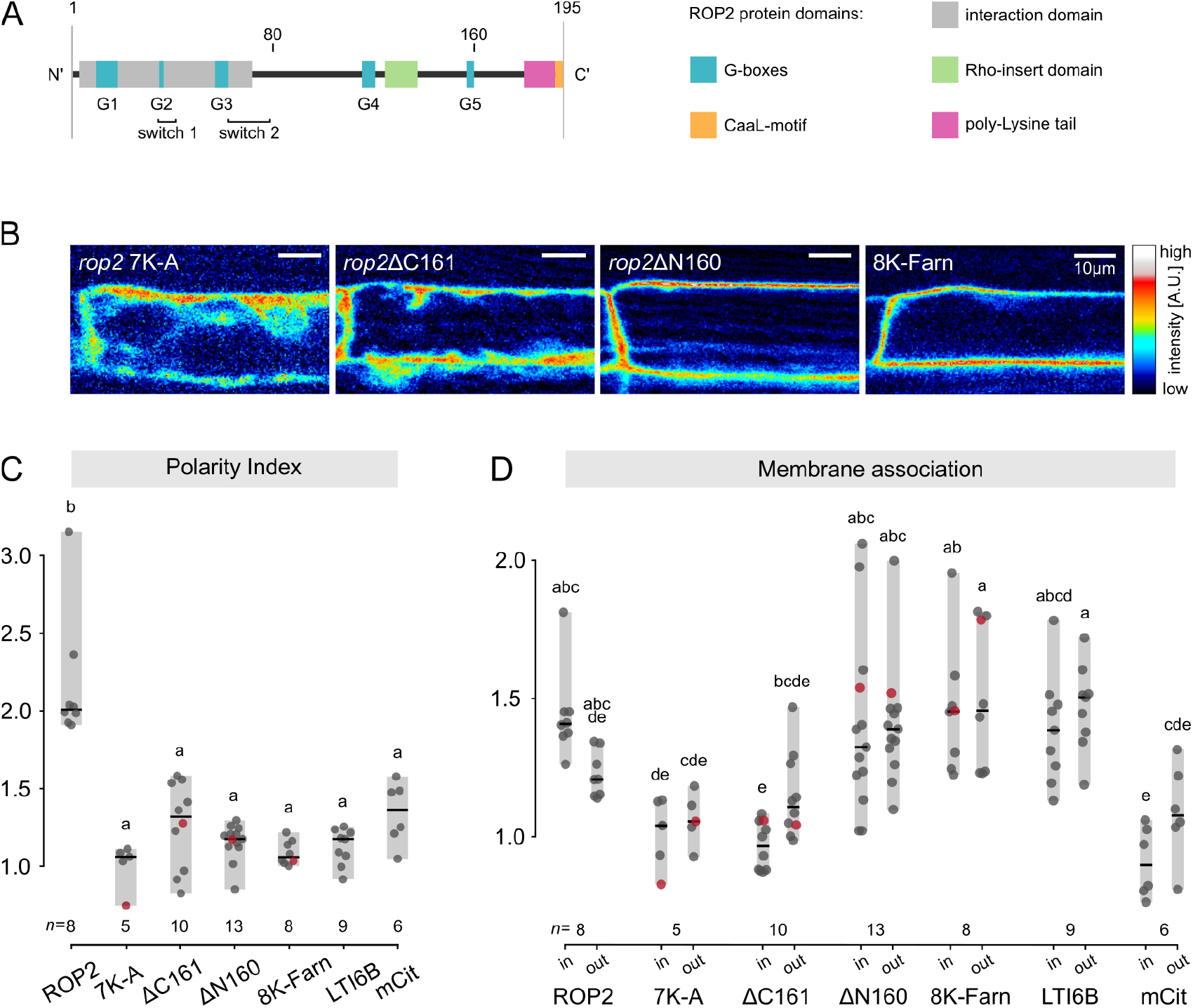
Association with the membrane involving electrostatic interactions with anionic lipids is a prerequisite for ROP2 polarization. (A) schematic representation of the primary protein structure of ROP2 depicting different functional and structural domains (B) Micrographs of trichoblast of cell stage -1 from plants expressing different ROP2 truncation constructs - ROP2 with a mutated polybasic tail (*rop2* 7K-A), ROP2 without the first 160 amino acids (*rop2*ΔN160), ROP2 without the C-terminal anchor (*rop2*ΔC161) and a farnesylated poly-lysine membrane anchor (8K-Farn) Scale bars represent 10 µm. (C) Polarity index and (D) membrane association of the in (B) depicted ROP2 truncation variants as well as ROP2, the 8K-Farn membrane anchor, GFP-LTI6B and mCit, measured inside (in) and outside (out) of the RHID of trichoblasts of cell stage -1. Note that the data for ROP2, LTI6B and mCit is the same as in Fig.4 and is shown for comparability. Centre lines represent median values, gray boxes represent the data range, n indicates the number of cells measured and letters represent the result of an ANOVA-Tukey test (significance value = 0.01; same letters indicate no significant difference). Red data points indicate data derived from the individuals shown in panel B.

**Fig. S5.**
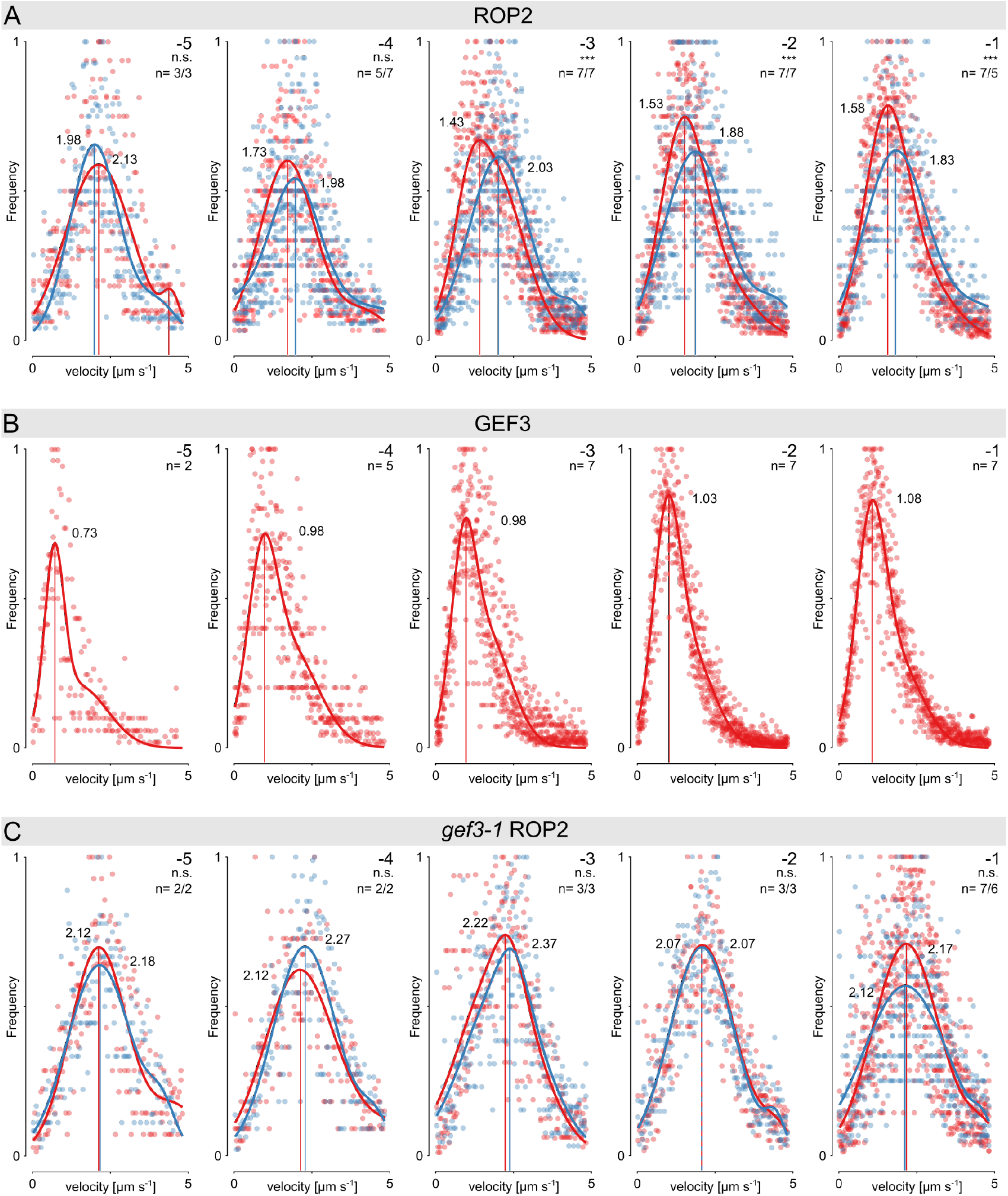
Protein mobility over the course of RHID development. Normalized distribution of particle velocities determined by ToBF for mCit-ROP2(A), mCit-GEF3 (B) and for mCit-ROP2 in the *gef3-1* mutant background (C) for the cell stages -5 to -1, measured inside (red) and outside (blue) of the RHID. p-values indicate the statistical difference between the distribution of velocities in- and outside the RHID determined by two-way ANOVA: n.s. = p-value >0.05; * = p-value < 0.05; ** = p-value < 0.01; *** = p-value < 0.001. Curves represent multi-gaussian fits of the pooled data sets, local maxima are indicated by vertical lines; n represents the number of biological replicates and is indicated for inside/outside the RHID in panel (A) and (C).

**Fig. S6.**
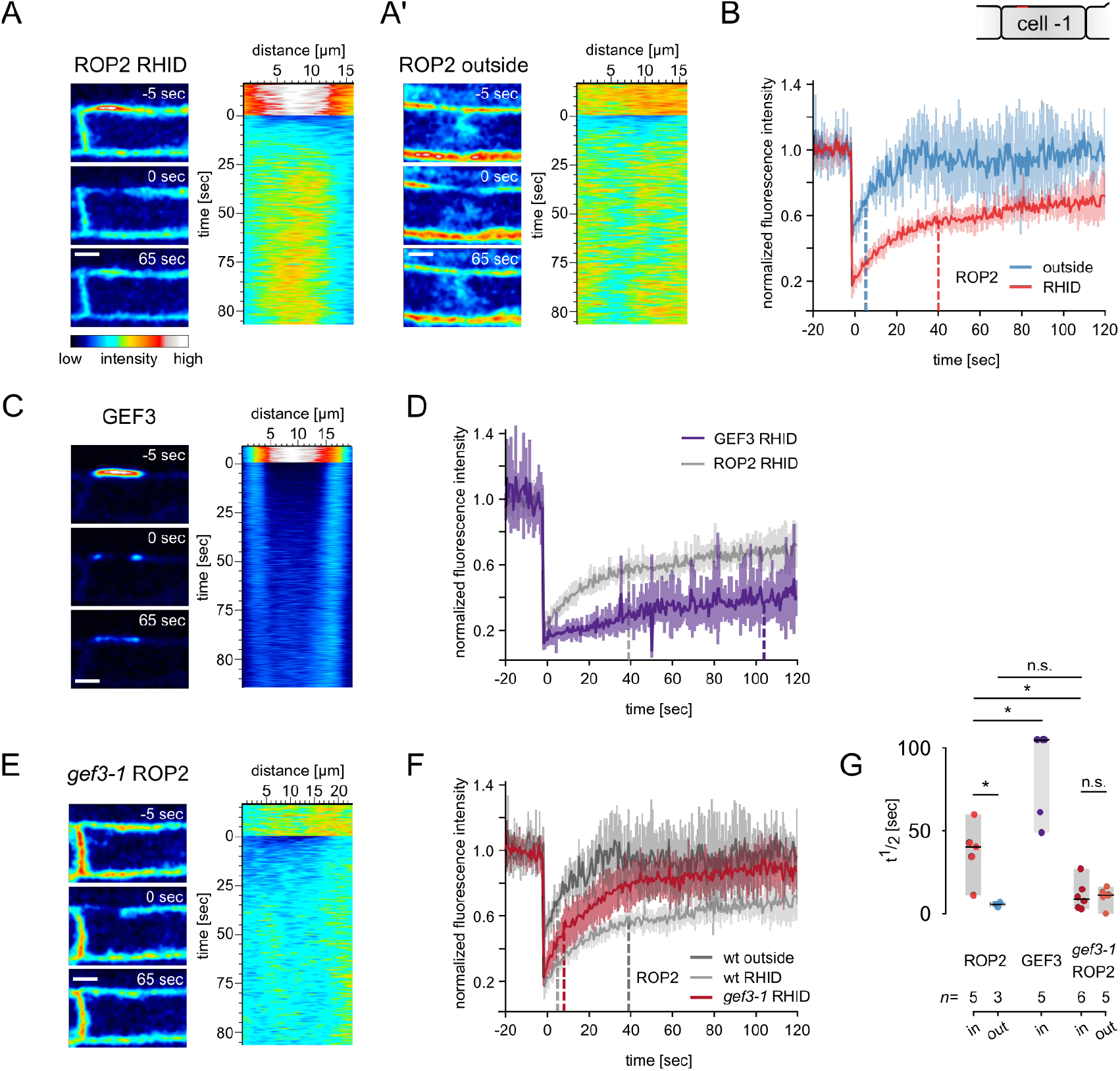
Immobilization of ROP2 in the RHID is dependent on the presence of GEF3. Fluorescent recovery after photobleaching (FRAP) measurements of ROP2 inside (A) and outside (A’) the RHID, of GEF3 (C) and of ROP2 in the *gef3-1* mutant background (E). Mean recovery-curve of the respective experiments for: ROP2 in- and outside the RHID (B) of GEF3 at the RHID (D) and of ROP2 in the *gef3-1* mutant background (F); dashed lines indicate the mean half time recovery (t½) value; data for ROP2 in D and F resemble the data in panel B and are shown for comparability. (G) Quantification of the half time recovery (t½) of ROP2, GEF3 and *gef3-1* ROP2 in- and outside the RHID; Centre lines represent median values, gray boxes represent the data range, n indicates the number of cells measured; p-value determined by Student’s t-test: n.s. = p-value >0.05; * = p-value < 0.05. All measurements were performed in cell stage -1. For each protein micrographs of single time points prior to (−5 sec), directly after (0 sec) and in the middle of the recorded time (65 sec), as well as a kymograph drawn along a line spanning the bleached region are shown; scale bars represent 10 µm.

**Fig. S7.**
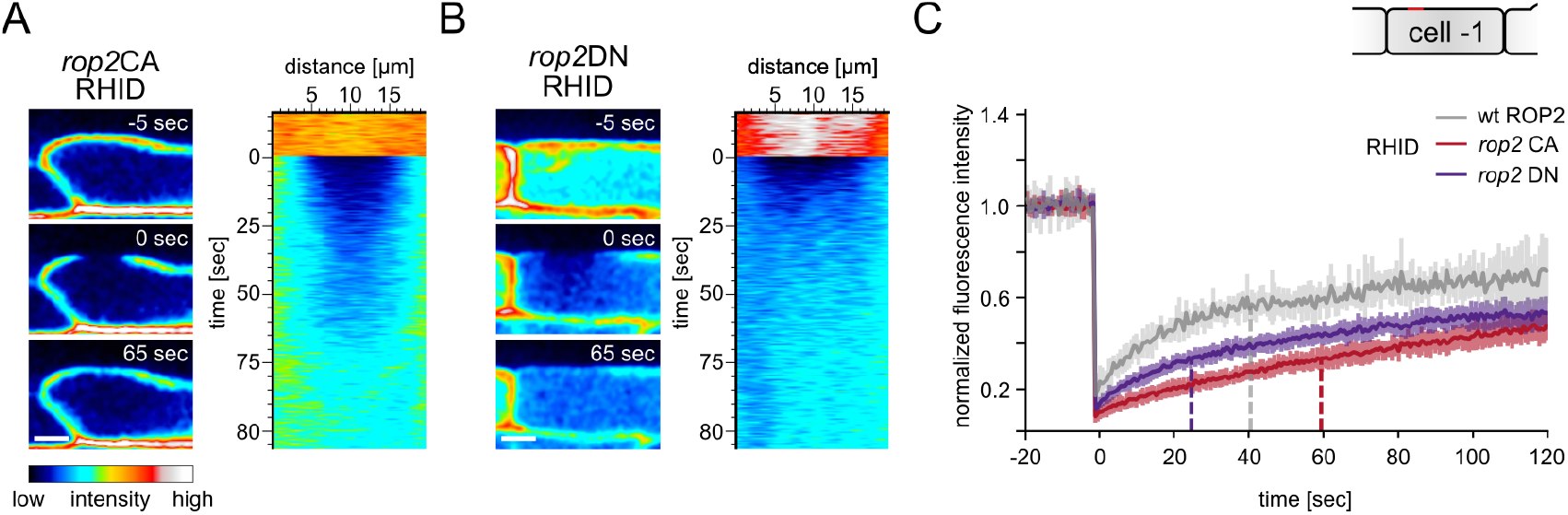
The activity state of ROP2 impacts on ROP2 protein mobility. Fluorescent recovery after photobleaching (FRAP) measurements of *rop2*CA (A) and *rop2*DN (B). (C) Mean recovery-curve of the respective experiments; dashed lines indicate the mean half time recovery (t½) value; data for ROP2 resembles the data in Fig.S6 and are shown for comparability. All measurements were performed in cell stage -1. For each protein micrographs of single time points prior to (−5 sec), directly after (0 sec) and in the middle of the recorded time (65 sec), as well as a kymograph drawn along a line spanning the bleached region are shown; scale bars represent 10 µm.

## References

Claire Grierson, Erik Nielsen, Tijs Ketelaarc, and John Schiefelbein. Root Hairs. The Arabidopsis Book 12:e0172, 2014. ISSN 1543-8120. doi: 10.1199/tab.0172.

Philipp Denninger, Anna Reichelt, Vanessa A.F. Schmidt, Dietmar G Mehlhorn, Lisa Y Asseck, Claire E Stanley, Nana F Keinath, Jan Felix Evers, Christopher Grefen, and Guido Grossmann. Distinct RopGEFs Successively Drive Polarization and Outgrowth of Root Hairs. Current Biology, 29(11):1854–1865, 2019. doi: 10.1016/j.cub.2019.04.059.

Gil Feiguelman, Ying Fu, and Shaul Yalovsky. ROP GTPases structure-function and signaling pathways, jan 2018. ISSN 15322548.

Zhi-Liang Zheng and Zhenbiao Yang. The Rop GTPase: an emerging signaling switch in plants. Plant Molecular Biology 2000 44:1, 44(1):1–9, 2000. ISSN 1573-5028. doi: 10.1023/A:1006402628948.

Rachel J. Carol, Seiji Takeda, Paul Linstead, Marcus C. Durrant, Hana Kakesova, Paul Derbyshire, Sinéad Drea, Viktor Zarsky, and Liam Dolan. A RhoGDP dissociation inhibitor spatially regulates growth in root hair cells. Nature, 438(7070):1013–1016, dec 2005. ISSN 14764687. doi: 10.1038/nature04198.

Mark A Jones, Jun-Jiang Shen, Ying Fu, Hai Li, Zhenbiao Yang, and Claire S Grierson. The Arabidopsis Rop2 GTPase is a positive regulator of both root hair initiation and tip growth. The Plant cell, 14(4):763–76, apr 2002. ISSN 1040-4651.

Catherine A. Konopka and Sebastian Y. Bednarek. Variable-angle epifluorescence microscopy: a new way to look at protein dynamics in the plant cell cortex. The Plant Journal, 53(1): 186–196, jan 2008. ISSN 09607412. doi: 10.1111/j.1365-313X.2007.03306.x.

Robert M. Dickson, Andrew B. Cubitt, Roger Y. Tsien, and W. E. Moerner. On/off blinking and switching behaviour of single molecules of green fluorescent protein. Nature, 388(6640): 355–358, jul 1997. ISSN 0028-0836. doi: 10.1038/41048.

Jonas Fölling, Mariano Bossi, Hannes Bock, Rebecca Medda, Christian A Wurm, Birka Hein, Stefan Jakobs, Christian Eggeling, and Stefan W Hell. Fluorescence nanoscopy by groundstate depletion and single-molecule return. Nature Methods, 5(11):943–945, nov 2008. ISSN 1548-7091. doi: 10.1038/nmeth.1257.

Jan Vogelsang, Christian Steinhauer, Carsten Forthmann, Ingo H. Stein, Britta Person-Skegro, Thorben Cordes, and Philip Tinnefeld. Make them Blink: Probes for Super-Resolution Microscopy. ChemPhysChem, 11(12):2475–2490, aug 2010. ISSN 1439-7641. doi: 10.1002/CPHC.201000189.

Julia Riedl, Alvaro H. Crevenna, Kai Kessenbrock, Jerry Haochen Yu, Dorothee Neukirchen, Michal Bista, Frank Bradke, Dieter Jenne, Tad A. Holak, Zena Werb, Michael Sixt, and Roland Wedlich-Soldner. Lifeact: A versatile marker to visualize F-actin. Nature Methods, 5(7):605–607, jul 2008. ISSN 15487091. doi: 10.1038/nmeth.1220.

Nils Gustafsson, Siân Culley, George Ashdown, Dylan M Owen, Pedro Matos Pereira, and Ricardo Henriques. Fast live-cell conventional fluorophore nanoscopy with ImageJ through super-resolution radial fluctuations. Nature Communications, 7:12471, aug 2016.

Matthieu Pierre Platre, Lise C. Noack, Mehdi Doumane, Vincent Bayle, Mathilde Laetitia Audrey Simon, Lilly Maneta-Peyret, Laetitia Fouillen, Thomas Stanislas, Laia Armengot, Přemysl Pejchar Marie Cécile Caillaud, Martin Potocký, Alenka Čopič, Patrick Moreau, and Yvon Jaillais. A Combinatorial Lipid Code Shapes the Electrostatic Landscape of Plant Endomembranes. Developmental Cell, 45(4):465–480.e11, 2018. ISSN 18781551. doi: 10.1016/j.devcel.2018.04.011.

Geert van den Bogaart, Karsten Meyenberg, H Jelger Risselada, Hayder Amin, Katrin I Willig, Barbara E Hubrich, Markus Dier, Stefan W Hell, Helmut Grubmüller, Ulf Diederichsen, and Reinhard Jahn. Membrane protein sequestering by ionic protein-lipid interactions. Nature, 479(7374):552–5, nov 2011. ISSN 1476-4687. doi: 10.1038/nature10545.

Benedikt Kost, Emmanuel Lemichez, Pius Spielhofer, Yan Hong, Kimberly Tolias, Christopher Carpenter, and Nam Hai Chua. Rac homologues and compartmentalized phosphatidylinositol 4,5-bisphosphate act in a common pathway to regulate polar pollen tube growth. Journal of Cell Biology, 145(2):317–330, jun 1999. ISSN 00219525. doi: 10.1083/jcb.145.2.317.

Markus Braun, Frantisek Baluška, Matthias von Witsch, and Diedrik Menzel. Redistribution of actin, profilin and phosphatidylinositol-4,5-bisphosphate in growing and maturing root hairs. Planta 1999 209:4, 209(4):435–443, oct 1999. ISSN 1432-2048. doi: 10.1007/S004250050746.

Hiroaki Kusano, Christa Testerink, Joop E M Vermeer, Tomohiko Tsuge, Hiroaki Shimada, Atsuhiro Oka, Teun Munnik, and Takashi Aoyama. The Arabidopsis Phosphatidylinositol Phosphate 5-Kinase PIP5K3 is a key regulator of root hair tip growth. The Plant cell, 20(2):367–380, 2008. ISSN 1040-4651. doi: 10.1105/tpc.107.056119.

Amelie Mendrinna and Staffan Persson. Root hair growth: It’s a one way street. F1000Prime Reports, 7, feb 2015. ISSN 20517599. doi: 10.12703/P7-23.

Lise C. Noack and Yvon Jaillais. Functions of Anionic Lipids in Plants. Annual Review of Plant Biology, 71(1):71–102, apr 2020. ISSN 1543-5008. doi: 10.1146/annurev-arplant-081519-035910.

Matthieu Pierre Platre, Vincent Bayle, Laia Armengot, Joseph Bareille, Maria del Mar Marquès-Bueno, Audrey Creff, Lilly Maneta-Peyret, Jean Bernard Fiche, Marcelo Nollmann, Christine Miège, Patrick Moreau, Alexandre Martinière, and Yvon Jaillais. Developmental control of plant Rho GTPase nano-organization by the lipid phosphatidylserine. Science, 364(6435): 57–62, apr 2019. ISSN 10959203. doi: 10.1126/science.aav9959.

Mathilde Laetitia Audrey Simon, Matthieu Pierre Platre, Sonia Assil, Ringo van Wijk, William Yawei Chen, Joanne Chory, Marlène Dreux, Teun Munnik, and Yvon Jaillais. A multi-colour/multi-affinity marker set to visualize phosphoinositide dynamics in Arabidopsis. The Plant journal: for cell and molecular biology, 77(2):322–37, jan 2014. ISSN 1365-313X. doi: 10.1111/tpj.12358.

Mathilde Laetitia Audrey Simon, Matthieu Pierre Platre, Maria Mar Marquès-Bueno, Laia Armengot, Thomas Stanislas, Vincent Bayle, Marie-Cécile Caillaud, and Yvon Jaillais. A PtdIns(4)P-driven electrostatic field controls cell membrane identity and signalling in plants. Nature Plants, 2(7):16089, jul 2016. doi: 10.1038/nplants.2016.89.

S. R. Cutler, D. W. Ehrhardt, J. S. Griffitts, and C. R. Somerville. Random GFP::cDNA fusions enable visualization of subcellular structures in cells of Arabidopsis at a high frequency. Proceedings of the National Academy of Sciences, 97(7):3718–3723, mar 2000. ISSN 0027-8424. doi: 10.1073/pnas.97.7.3718.

Yoji Kawano, Takako Kaneko-Kawano, and Ko Shimamoto. Rho family GTPase-dependent immunity in plants and animals. Frontiers in Plant Science, 5:522, oct 2014. ISSN 1664-462X. doi: 10.3389/fpls.2014.00522.

A J Molendijk, F Bischoff, C S Rajendrakumar, J Friml, M Braun, S Gilroy, and K Palme. Arabidopsis thaliana Rop GTPases are localized to tips of root hairs and control polar growth. The EMBO journal, 20(11):2779–88, jun 2001. ISSN 0261-4189. doi: 10.1093/emboj/20.11.2779.

Suliana Manley, Jennifer M. Gillette, George H. Patterson, Hari Shroff, Harald F. Hess, Eric Betzig, and Jennifer Lippincott-Schwartz. High-density mapping of single-molecule trajectories with photoactivated localization microscopy. Nature Methods, 5(2):155–157, feb 2008. ISSN 15487091. doi: 10.1038/nmeth.1176.

Vincent Bayle, Jean Bernard Fiche, Claire Burny, Matthieu Pierre Platre, Marcelo Nollmann, Alexandre Martinière, and Yvon Jaillais. Single-particle tracking photoactivated localization microscopy of membrane proteins in living plant tissues. Nature Protocols, 16(3):1600–1628, mar 2021. ISSN 17502799. doi: 10.1038/s41596-020-00471-4.

Mike Heilemann, Sebastian van de Linde, Mark Schüttpelz, Robert Kasper, Britta Seefeldt, Anindita Mukherjee, Philip Tinnefeld, and Markus Sauer. Subdiffraction-Resolution Fluorescence Imaging with Conventional Fluorescent Probes. Angewandte Chemie International Edition, 47(33):6172–6176, aug 2008. ISSN 14337851. doi: 10.1002/anie.200802376.

Jürgen Kleine-Vehn, Krzysztof Wabnik, Alexandre Martinière, Łukasz Łangowski, Katrin Willig, Satoshi Naramoto, Johannes Leitner, Hirokazu Tanaka, Stefan Jakobs, Stéphanie Robert, Christian Luschnig, Willy Govaerts, Stefan W Hell, John Runions, and Jiří Friml. Recycling, clustering, and endocytosis jointly maintain PIN auxin carrier polarity at the plasma membrane. Molecular systems biology, 7(540):540, jan 2011. ISSN 1744-4292. doi: 10.1038/msb.2011.72.

Delphine Gendre, Anirban Baral, Xie Dang, Nicolas Esnay, Yohann Boutté, Thomas Stanislas, Thomas Vain, Stéphane Claverol, Anna Gustavsson, Deshu Lin, Markus Grebe, and Rishikesh P. Bhalerao. Rho-of-plant-activated root hair formation requires Arabidopsis YIP4a/b gene function. Development, 146(5):1–7, jan 2019. ISSN 1477-9129. doi: 10.1242/dev.168559.

Marta Fratini, Praveen Krishnamoorthy, Irene Stenzel, Mara Riechmann, Monique Matzner, Kirsten Bacia, Mareike Heilmann, and Ingo Heilmann. Plasma membrane nano-organization specifies phosphoinositide effects on Rho-GTPases and actin dynamics in tobacco pollen tubes. The Plant Cell, 33(3):642–670, may 2021. ISSN 1532-298X. doi: 10.1093/plcell/koaa035.

Michael Glotzer and Anthony A. Hyman. Cell Polarity: The importance of being polar. Current Biology, 5(10):1102–1105, oct 1995. ISSN 09609822. doi: 10.1016/S0960-9822(95)00221-1.

Antje Berken and Alfred Wittinghofer. Structure and function of Rho-type molecular switches in plants, mar 2008. ISSN 09819428.

Roland Wedlich-Soldner and Rong Li. Spontaneous cell polarization: undermining determinism. Nature cell biology, 5(4):267–70, apr 2003. ISSN 1465-7392. doi: 10.1038/ncb0403-267.

Jian-geng Chiou, Mohan K. Balasubramanian, and Daniel J. Lew. Cell Polarity in Yeast. Annual Review of Cell and Developmental Biology, 33(1):77–101, oct 2017. ISSN 1081-0706. doi: 10.1146/annurev-cellbio-100616-060856.

Urs Fischer, Yoshihisa Ikeda, Karin Ljung, Olivier Serralbo, Manoj Singh, Renze Heidstra, Klaus Palme, Ben Scheres, and Markus Grebe. Vectorial Information for Arabidopsis Planar Polarity Is Mediated by Combined AUX1, EIN2, and GNOM Activity. Current Biology, 16(21):2143–2149, nov 2006. ISSN 09609822. doi: 10.1016/j.cub.2006.08.091.

Robert J. H. Payne and Claire Suzanne Grierson. A Theoretical Model for ROP Localisation by Auxin in Arabidopsis Root Hair Cells. PLoS ONE, 4(12):e8337, dec 2009. ISSN 1932-6203. doi: 10.1371/journal.pone.0008337.

Ivan Kulich, Frank Vogler, Andrea Bleckmann, Philipp Cyprys, Maria Lindemeier, Ingrid Fuchs, Laura Krassini, Thomas Schubert, Jens Steinbrenner, Jim Beynon, Pascal Falter-Braun, Gernot Längst, Thomas Dresselhaus, and Stefanie Sprunck. ARMADILLO REPEAT ONLY proteins confine Rho GTPase signalling to polar growth sites. Nature Plants, 6(10):1275–1288, oct 2020. ISSN 20550278. doi: 10.1038/s41477-020-00781-1.

Julien Gronnier, Patricia Gerbeau-Pissot, Véronique Germain, Sébastien Mongrand, and Françoise Simon-Plas. Divide and Rule: Plant Plasma Membrane Organization. Trends in Plant Science, 23(10):899–917, oct 2018. ISSN 1360-1385. doi: 10.1016/J.TPLANTS.2018.07.007.

Athanasios Lampropoulos, Zoran Sutikovic, Christian Wenzl, Ira Maegele, Jan U. Lohmann, and Joachim Forner. GreenGate - A Novel, Versatile, and Efficient Cloning System for Plant Transgenesis. PLoS ONE, 8(12):e83043, dec 2013. ISSN 1932-6203. doi: 10.1371/journal.pone.0083043.

Nathan C. Shaner, Gerard G. Lambert, Andrew Chammas, Yuhui Ni, Paula J. Cranfill, Michelle A. Baird, Brittney R. Sell, John R. Allen, Richard N. Day, Maria Israelsson, Michael W. Davidson, and Jiwu Wang. A bright monomeric green fluorescent protein derived from Branchiostoma lanceolatum. Nature Methods, 10(5):407–409, 2013. ISSN 15487091. doi: 10.1038/nmeth.2413.

Grigorios Koulouras, Andreas Panagopoulos, Maria A. Rapsomaniki, Nickolaos N. Giakoumakis, Stavros Taraviras, and Zoi Lygerou. EasyFRAP-web: A web-based tool for the analysis of fluorescence recovery after photobleaching data. Nucleic Acids Research, 2018. ISSN 13624962. doi: 10.1093/nar/gky508.

Dietmar G. Mehlhorn, Niklas Wellmeroth, Kenneth W. Berendzen, and Christopher Grefen. 2in1 Vectors Improve In Planta BiFC and FRET Analyses. In Methods in Molecular Biology, volume 1691, pages 139–158. Humana Press, New York, NY, 2018. ISBN 978-1-4939-7388-0. doi: https://doi.org/10.1007/978-1-4939-7389-7_11.

Johannes Schindelin, Ignacio Arganda-Carreras, Erwin Frise, Verena Kaynig, Mark Longair, Tobias Pietzsch, Stephan Preibisch, Curtis Rueden, Stephan Saalfeld, Benjamin Schmid, Jean-Yves Tinevez, Daniel James White, Volker Hartenstein, Kevin Eliceiri, Pavel Tomancak, and Albert Cardona. Fiji: an open-source platform for biological-image analysis. Nature methods, 9(7):676–82, jul 2012. ISSN 1548-7105. doi: 10.1038/nmeth.2019.

Jean-Yves Tinevez, Nick Perry, Johannes Schindelin, Genevieve M. Hoopes, Gregory D. Reynolds, Emmanuel Laplantine, Sebastian Y. Bednarek, Spencer L. Shorte, and Kevin W. Eliceiri. TrackMate: An open and extensible platform for single-particle tracking. Methods, 115:80–90, feb 2017. ISSN 1046-2023. doi: 10.1016/J.YMETH.2016.09.016.

Hadley Wickham. ggplot2: Elegant Graphics for Data Analysis. Springer-Verlag New York, 2016. ISBN 978-3-319-24277-4.

Jeffrey B. Arnold. ggthemes: Extra Themes, Scales and Geoms for ‘ggplot2’, 2019.

John Fox and Sanford Weisberg. An R Companion to Applied Regression. Sage, Thousand Oaks CA, third edition, 2019.

RStudio Team. RStudio: Integrated Development for R. RStudio, 2020.

